# An inherited mtDNA mutation remodels inflammatory cytokine responses in macrophages and *in vivo*

**DOI:** 10.1101/2025.01.05.631298

**Authors:** Eloïse Marques, Stephen P. Burr, Alva M. Casey, Richard J. Stopforth, Chak Shun Yu, Keira Turner, Dane M. Wolf, Marisa Dilucca, Victoria J. Tyrrell, Robbin Kramer, Yamini M. Kanse, Chris A. Powell, James B. Stewart, Michael P. Murphy, Michal Minczuk, Valerie B. O’Donnell, Clare E. Bryant, Patrick F. Chinnery, Arthur Kaser, Alexander von Kriegsheim, Dylan G. Ryan

## Abstract

Impaired mitochondrial bioenergetics in macrophages can drive hyperinflammatory cytokine responses^1–6^, but whether this may also be caused by inherited mtDNA mutations is unknown. Here, we address this question using a multi-omic approach that integrates super-resolution imaging and metabolic analyses to profile macrophages from a mouse model of mitochondrial disease arising from a heteroplasmic mutation (m.5019A>G) in the mitochondrial tRNA for alanine^7^. These m.5019A>G macrophages exhibit defects in respiratory chain complexes and oxidative phosphorylation (OxPhos) due to decreased intra-mitochondrial translation. To adapt to this metabolic stress, mitochondrial fusion, reductive glutamine metabolism, and aerobic glycolysis are all increased. Upon inflammatory activation, type I interferon (IFN-I) release is enhanced, while the production of pro-inflammatory cytokines and oxylipins are restrained in m.5019A>G macrophages. Finally, an *in vivo* endotoxemia model using m.5019A>G mice reveal elevated IFN-I levels and sickness behaviour. In conclusion, our study identifies an unexpected imbalance in innate immune signalling in response to a pathogenic mtDNA mutation, with important implications for the progression of pathology in patients with mtDNA diseases^8^.

## Main

Mitochondria are intracellular organelles that act as a nexus for the integration of anabolic and catabolic pathways essential to eukaryotic life^9–11^. They play a central role in cellular bioenergetics as the main producers of ATP via oxidative phosphorylation (OxPhos), as well as in the supply of intermediates for the synthesis of all major biological macromolecule subtypes. This organelle contains its own circular chromosome of approximately 16.5 kb, termed mitochondrial DNA (mtDNA)^12^. Importantly, mtDNA encodes 37 genes including 2 ribosomal RNAs, 22 tRNAs and 13 protein subunits that are critical components of the mitochondrial respiratory chain, while most other genes encoding mitochondrial proteins are in the nucleus.

Multiple copies of mtDNA are found per cell depending on cellular energetic demands and are uniparentally inherited through the maternal germline^12,13^. Inherited and somatic mutations in mtDNA give rise to heteroplasmy, the coexistence of one or more variants of mtDNA within a cell^12,14^. Somatic heteroplasmic single nucleotide variants arise throughout the human lifespan and accumulate sharply after 70 years of age^14^. Since their discovery approximately 30 years ago, inherited mtDNA mutations have emerged as a key driver of primary mitochondrial disease, a group of genetic disorders characterised by impairments in mitochondrial bioenergetics affecting ∼1 in 4,300 of the human population^13,15^. The largest proportion of heteroplasmic mtDNA mutations occur in genes encoding mitochondrial tRNAs and disrupt intra-mitochondrial translation^13^. Recurrent microbial infection, sepsis and systemic inflammatory response syndrome (SIRS) are commonly observed in mitochondrial disease patients and a major cause of morbidity and mortality^8,16–21^. Despite this, we do not understand how pathogenic mtDNA mutations impact the innate immune system.

Macrophages are essential cells of the innate immune system^22^. Metabolic rewiring underlies the essential functional plasticity of these cells by supporting pathogen clearance, intra-and inter-cellular communication, and the resolution of inflammation^9,23,24^. Mitochondria are central to this metabolic rewiring, serving as vital signalling hubs for the execution of macrophage effector functions following activation^9,25,26^. Previous reports studying mouse models of mitochondrial dysfunction have shown that macrophages drive pathological hyperinflammatory responses^1–4,27^. These models have typically relied on profiling *Polg*^D257A^ mutant or *Ndufs4*^-/-^ (complex I subunit) macrophages, which are nuclear-encoded mitochondrial proteins often presenting with severe phenotypes^1–4,6,27^. The *Polg*^D257A^ mutation is found in the catalytic subunit of the mtDNA polymerase and impairs proof-reading^28^. This loss of proof-reading activity leads to the damage and depletion of mtDNA. However, this does not replicate inherited pathogenic mtDNA mutations found in patients with mtDNA disease. Therefore, it is unclear how heritable pathogenic mtDNA mutations impact inflammatory macrophage activation and inflammation *in vivo*.

To address this outstanding question, we used the m.5019A>G mouse, a recently developed heteroplasmic mtDNA mutation model^7^. Here, we find that primary macrophages with the m.5019A>G mutation exhibit impaired mitochondrial respiration due to combined disruption of respiratory chain complex I (CI), CIII and CIV, which results in a stress-induced mitochondrial hyperfusion (SIMH) phenotype with disrupted cristae architecture. To compensate for these respiratory chain defects, macrophages engage in reductive glutamine metabolism and aerobic glycolysis. However, this disrupts inflammatory macrophage activation increasing IFN-I release and inducible nitric oxide synthase (iNOS)-dependent nitric oxide (NO) production, while limiting pro-inflammatory cytokine and oxylipin production. IFN-I release is also elevated in the serum of m.5019A>G mice following a lipopolysaccharide (LPS) challenge *in vivo*. Together, our data suggests that heteroplasmic mtDNA mutations perturb innate immune responses *ex vivo* and *in vivo*, which may have implications for mtDNA disease patients.

## Results

### Characterisation of m.5019A>G macrophages

The m.5019A>G mouse contains a point mutation in the mitochondrial tRNA^Ala^ gene (mt-*Ta*) (**Fig. 1a**), which occurs in the acceptor stem of mt-*Ta* and prevents charging with its cognate amino acid, alanine^7^. Heteroplasmy proportion was determined by pyrosequencing of DNA extracted from ear skin biopsies obtained at weaning. Only mice with a high proportion of the m.5019A>G mutation (between 70% and 87%) were used in this study. Pyrosequencing was also performed on bone marrow pre-differentiation and primary bone marrow-derived macrophages (BMDMs) post-differentiation, confirming high mutational burdens that remained stable throughout the differentiation process (**Fig. 1b**). There was no negative impact on macrophage differentiation, as assessed by cell surface expression of F4/80 and proteomic measurements of F4/80, CD11b and CD11c (**Extended data Fig. 1a, b**). However, there was a significant decrease in MHC class II protein levels (**Extended data Fig. 1c**). mtDNA copy number was similar in non-stimulated (non-stim) WT and m.5019A>G macrophages, whereas there was an increase in m.5019A>G macrophages following stimulation with LPS from gram-negative bacterial cell membranes, a potent pro-inflammatory agent (**Fig. 1c and extended data Fig. 1d**). To determine whether the m.5019A>G mutation had an impact on mitochondrial translation in macrophages, we performed ^35^S-methionine labelling of mitochondrial proteins (**extended data Fig. 1e**). ^35^S-methionine incorporation was significantly reduced in non-stim m.5019A>G macrophages compared with WT macrophages, while complex IV (CIV)-subunit MT-CO1 was also reduced (**Extended data Fig. 1e, f**). There was no decrease in the transcript levels of *mt-Nd1* or *mt-Co3* (**Extended data Fig. 1g**), confirming the defect occurs at the level of mitochondrial translation.

**Figure 1.**
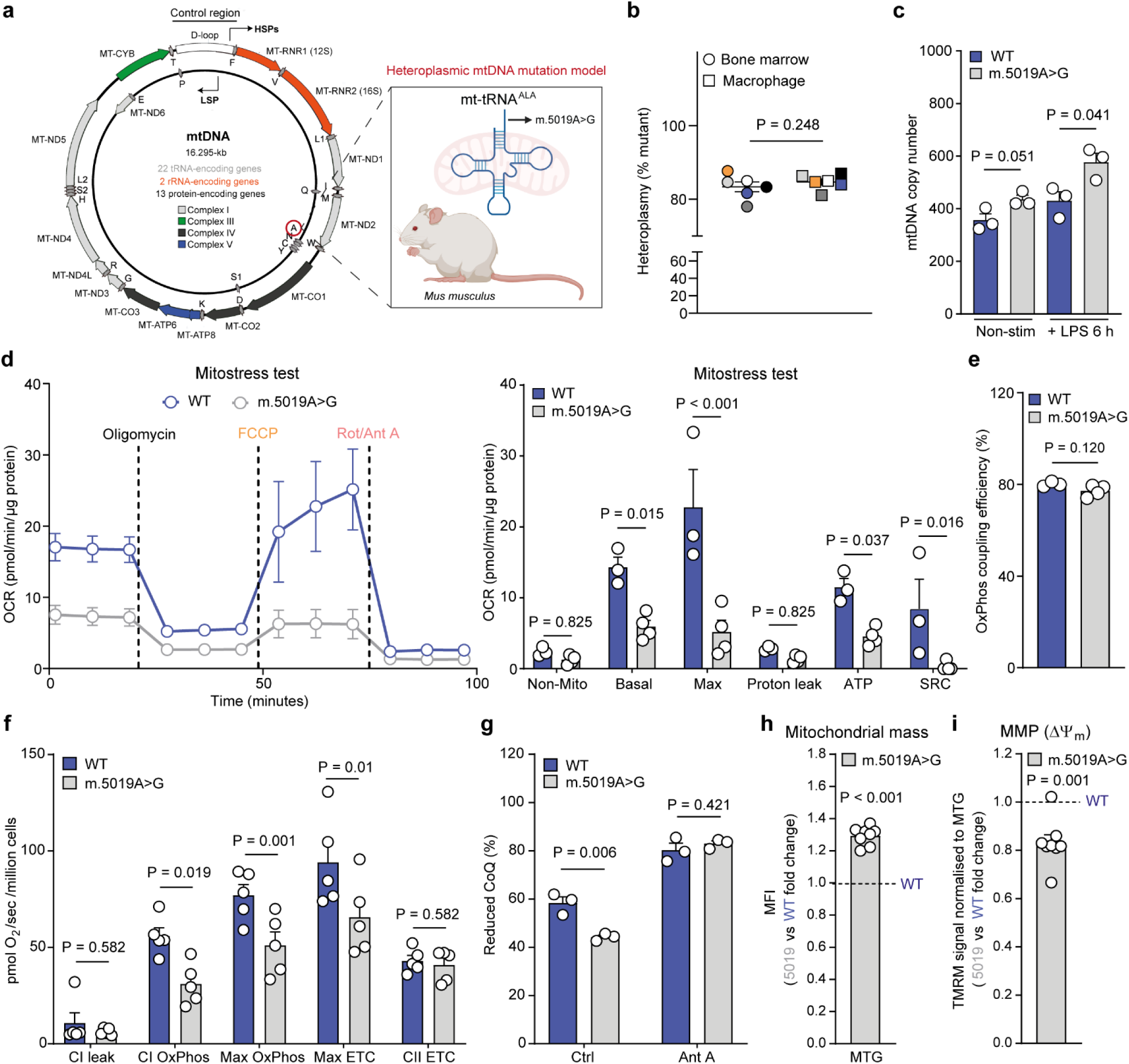
Reduced mitochondrial respiration in primary macrophages with a heteroplasmic m.5019A>G mtDNA mutation **a**, Schematic of m.5019A>G mt-*Ta* mutation model. **b**, Pyrosequencing results of bone marrow and bone marrow-derived macrophages (BMDMs) from m.5019A>G mice (*n* = 6). **c**, mtDNA copy number in non-stim and LPS-stimulated WT and m.5019A>G BMDMs (*n* = 3; LPS 6 h). **d-e**, Respirometry analysis using the Mitostress test in non-stim WT and m.5019A>G BMDMs (*n* = 3-4). **f**, Respirometry analysis of permeabilised non-stim WT and m.5019A>G BMDMs (*n* = 5). ETC = electron transfer capacity. **g**, CoQ redox measurements with or without Antimycin A (Ant A) in non-stim WT and m.5019A>G BMDMs (*n* = 3). **h**, Mitochondrial mass and (**i**) normalised mitochondrial membrane potential measurements in non-stim m.5019A>G vs WT BMDMs using MitoTracker Green (MTG) and tetramethyl rhodamine methyl ester (TMRM) (*n* = 8). Data are mean ± s.e.m. *P* values calculated using two-tailed Student’s t-test for two group comparisons or multiple unpaired or paired t-tests corrected for multiple comparisons using Holm-Sidak method.

In agreement with a mitochondrial translation defect, there was a significant impairment in basal respiration, spare respiratory capacity (SRC) and ATP-linked oxygen consumption in non-stim m.5019A>G macrophages (**Fig. 1d**). However, no difference in proton leak or OxPhos coupling efficiency was observed (**Fig. 1d, e**), which demonstrates that mitochondrial ATP synthesis can still occur albeit to a lesser extent. Further respirometry analysis in permeabilised cells revealed a significant reduction in CI-dependent respiration in non-stim m.5019A>G macrophages, which was not observed for CII (**Fig. 1f and extended data Fig. 1h**). Consistent with a decrease in CI-dependent respiration, Coenzyme Q (CoQ) was significantly more oxidised in non-stim m.5019A>G macrophages compared to WT (**Fig. 1g**). This impairment in respiration coincided with an increase in mitochondrial mass (**Fig. 1h**) and a reduction in mitochondrial membrane potential (MMP) (**Fig. 1i** and **extended Fig. 1i**), as determined by MitoTracker Green (MTG) and tetramethyl rhodamine methyl ester (TMRM) staining followed by flow cytometric analysis.

### Disrupted inflammatory activation in m.5019A>G macrophages

To determine the impact of the m.5019A>G mt-*Ta* mutation on inflammatory macrophage function, we applied three orthogonal approaches. We performed RNA sequencing to identify differentially expressed inflammatory genes (**Fig. 2a, b**), cytokine profiling of the cell culture medium (CCM) using Olink technology to identify differentially released cytokines and chemokines (**Fig. 2d**), and oxylipin profiling of the CCM with liquid chromatography tandem-mass spectrometry (LC-MS/MS) to identify differentially produced inflammatory lipid mediators (**Fig. 2e**) in LPS-stimulated WT and m.5019A>G macrophages. A robust increase in interferon-beta (IFN-β) gene expression (*Ifnb1*) was observed in LPS-stimulated m.5019A>G macrophages (**Fig. 2a**), consistent with previous reports of mitochondrial respiration inhibition in LPS-activated macrophages^29^ and the *Polg*^D257A^ mouse model of mitochondrial disease^1,3^. Overrepresentation analysis (ORA) using KEGG pathways indicated an enrichment in viral-associated inflammation (**Fig. 2b**) and increased IFN-β release was confirmed using an ELISA assay (**Fig. 2c**), which was more pronounced after 24 h of stimulation (**Extended data Fig. 2a**). In addition, we also identified a significant increase in immunostimulatory mtRNA and mtDNA (**Extended data Fig. 2b-d**) in the cytosol of m.5019A>G macrophages, which has previously been shown to activate the double stranded (dsRNA) sensors, RIG-I and MDA5, and the DNA sensor, cGAS, respectively, to drive IFN-β release in LPS-stimulated macrophages^1,29,30^.

**Figure 2.**
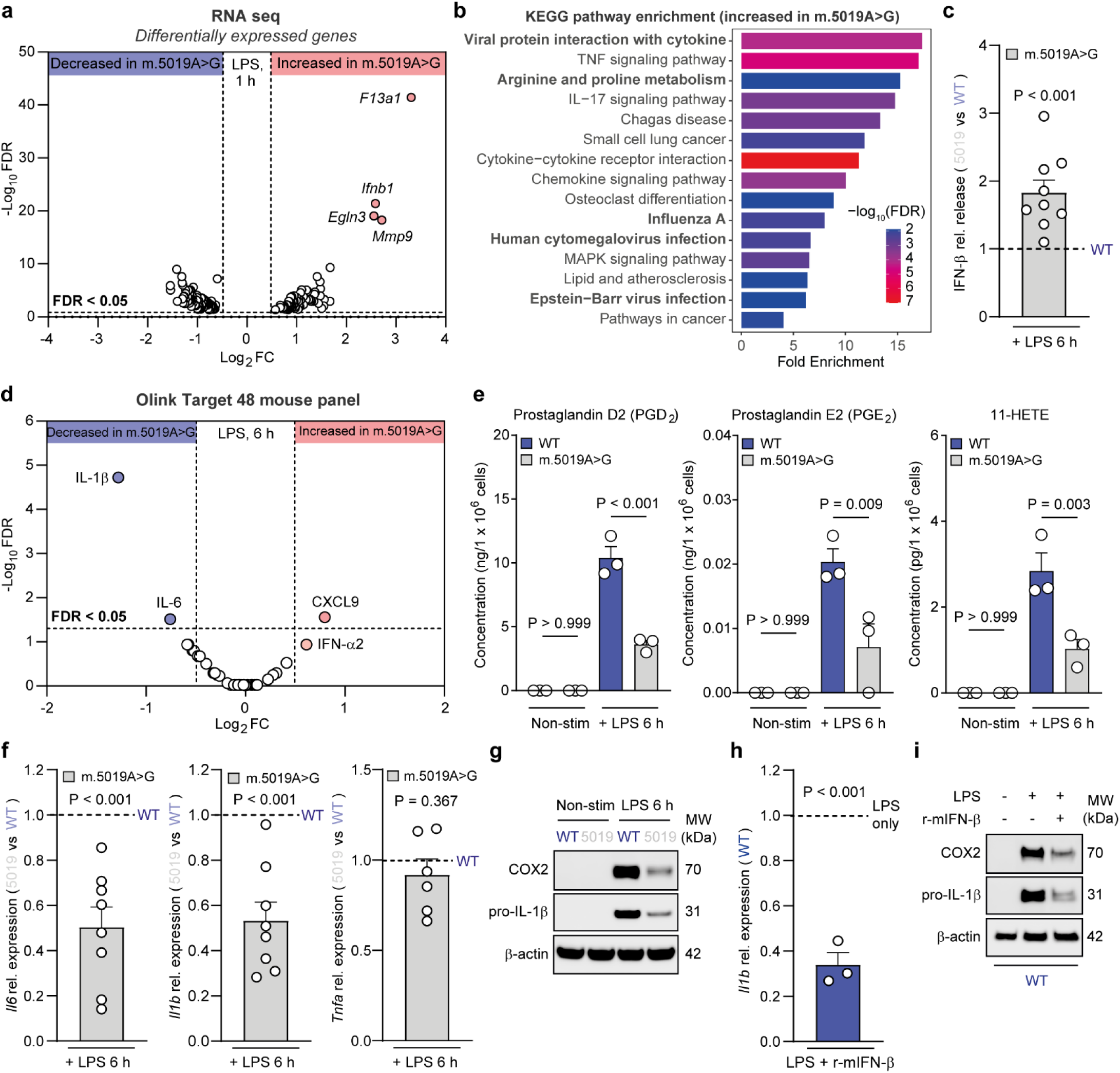
Inflammatory cytokine and oxylipin production are disrupted in m.5019A>G primary macrophages **a**, Volcano plot of differentially expressed genes from RNA sequencing (LPS 1 h) and (**b**) ORA analysis using KEGG pathway enrichment in genes increased in m.5019A>G vs WT BMDMs (n = 3). **c**, IFN-β release in LPS-stimulated m.5019A>G vs WT BMDMs (*n* = 9; LPS 6 h). **d**, Olink target T48 mouse cytokine and chemokine profiling of CCM in LPS-stimulated m.5019A>G vs WT BMDMs (*n* = 3; LPS 6 h). **e**, Oxylipin profiling of CCM in non-stim and LPS-stimulated m.5019A>G vs WT BMDMs (*n* = 3; LPS 6 h). **f**, *Il6*, *Il1b* and *Tnfa* expression in LPS-stimulated m.5019A>G vs WT BMDMs (*n* = 6-8; LPS 6 h). **g**, pro-IL-1β and COX2 protein levels in non-stim and LPS-stimulated WT and m.5019A>G BMDMs (*n* = 5; LPS 6 h). Representative blot shown. **h**, *Il1b* expression (*n* = 3; 6 h) and (**i**) pro-IL-1β and COX2 protein levels in LPS and LPS with r- mIFN-β−stimulated WT macrophages (*n* = 3; 6 h; representative blot shown). Data are mean ± s.e.m. *P* values calculated using two-tailed Student’s t-test for two group comparisons, multiple unpaired t-tests corrected for multiple comparisons using two-stage step-up method of Benjamini, Krieger and Yekutieli method, or one-way ANOVA corrected for multiple comparisons using Tukey method.

Interestingly, Olink profiling of the CCM identified a significant decrease in IL-1β and IL-6 levels (**Fig. 2d**). IFN-β is not included as part of the profiling, however, there was a non-significant increase in the type I IFN (IFN-I), IFN-α2, which is less abundant than IFN-β in macrophages. Furthermore, oxylipin profiling identified and quantified six oxylipins in total in the CCM at this timepoint (**Fig. 2e and extended data Fig. 2e**), with a significant reduction in inflammatory cyclooxygenase (COX) products prostaglandin D2 (PGD_2_), PGE_2_ and 11-HETE (**Fig. 2e**) but not COX-independent oxylipins (**Extended data Fig. 2e**) in m.5019A>G macrophages. Decreases in *Il1b*, *Il6* and the inducible COX isoform, COX-2 (*Ptgs2*), were validated using a combination of qPCR, Western blot and ELISA-based analyses (**Fig. 2f, g and extended data Fig. 2f**). In contrast, no significant differences were observed in *Tnfa* expression in m.5019A>G macrophages at this timepoint (**Fig. 2f**), although there was a trending increase in TNF-α release (**Extended data Fig. 2f**). IL-1β release was also significantly decreased following infection with the gram-negative bacterium *Salmonella typhimurium* (STM) in LPS-primed m.5019A>G macrophages, while no difference in cell death or bacterial burden was observed (**Extended data Fig. 2g**). This indicates that reduced IL-1β release arises from reduced *Il1b* expression and pro-IL-1β levels. IFN-I is reported to antagonise IL-1β production^29,31,32^. Consistent with these reports, co-treatment of WT macrophages with LPS and recombinant murine IFN-β (r-mIFN-β) significantly decreased both pro-IL-1β and COX2 levels when compared with LPS alone, phenocopying observations in LPS-stimulated m.5019A>G macrophages (**Fig. 2h, i**). This suggests that elevated IFN-I may be a contributor to the observed inflammatory phenotype in m.5019A>G macrophages *ex vivo*. As such, the m5019A>G mt-*Ta* mutation disrupts inflammatory macrophage activation, enhancing IFN-I release, while selectively restricting IL-1β, IL-6 and COX-dependent inflammatory oxylipins.

### Stress-induced mitochondrial fusion and disrupted cristae architecture in m.5019A>G macrophages

To understand this imbalance in the innate immune response, we next sought to determine the downstream consequences of the m.5019A>G mt-*Ta* mutation on mitochondrial biology. Severe mitochondrial defects often give rise to mitochondrial fission and swelling^33,34^. Given the substantial impairment in mitochondrial respiration in m.5019A>G macrophages, we assessed whether this affected mitochondrial morphology. Contrary to our expectations, staining of the inner mitochondrial membrane (IMM) intermembrane space protein, Cytochrome c (Cyt c), and the outer mitochondrial membrane protein (OMM), TOMM20, revealed an increase in mitochondrial length, junction points and junction points per mitochondrial network in non-stim m.5019A>G macrophages, which remained elongated following LPS stimulation (**Fig. 3a, b and extended data Fig. 3a, b**). This observation is akin to previous reports of SIMH, a pro-survival adaptation to mild metabolic stress^34,35^. LPS stimulation also promoted mitochondrial elongation in WT macrophages to a similar extent as that observed in non-stim m.5019A>G macrophages (**Fig. 3a, b**). To confirm this phenotype, we used super-resolution microscopy to examine mitochondrial morphology by staining for TOMM20 and ATP synthase (CV) (**Fig. 3c**, **extended data Fig. 3c-e**). This confirmed the mitochondrial elongation phenotype as determined by a significant decrease in mitochondrial oblate ellipticity and sphericity (**Fig. 3d** and **extended data Fig. 3c**) in both non-stim and LPS-stimulated m.5019A>G macrophages. Interestingly, a significant increase in ATP synthase puncta (**Extended data Fig. 3c**) was observed in m.5019A>G macrophages, likely in compensation for reduced respiratory rates and mitochondrial ATP synthesis. Finally, we used transmission electron microscopy (TEM) to assess mitochondrial architecture (**Fig. 3e and extended data Fig. 3e**). Non-stim WT macrophages presented with more punctate electron dense mitochondria, which elongated following LPS stimulation (**Fig. 3e, f and extended data Fig. 3e**). In contrast, m.5019A>G mitochondria had a greater proportion of low-density elongated mitochondria with disrupted cristae architecture (**Fig. 3e, f and extended data Fig. 3e**), consistent with a defect in both mitochondrial respiration and SIMH. This data shows that the m.5019A>G mt-*Ta* mutation in macrophages leads to a profound remodelling of the mitochondrial network. Disruption of mitochondrial cristae architecture has previously been shown to induce IFN-I signalling^36^, and is consistent with the increase in IFN-β release observed in m.5019A>G macrophages.

**Figure 3.**
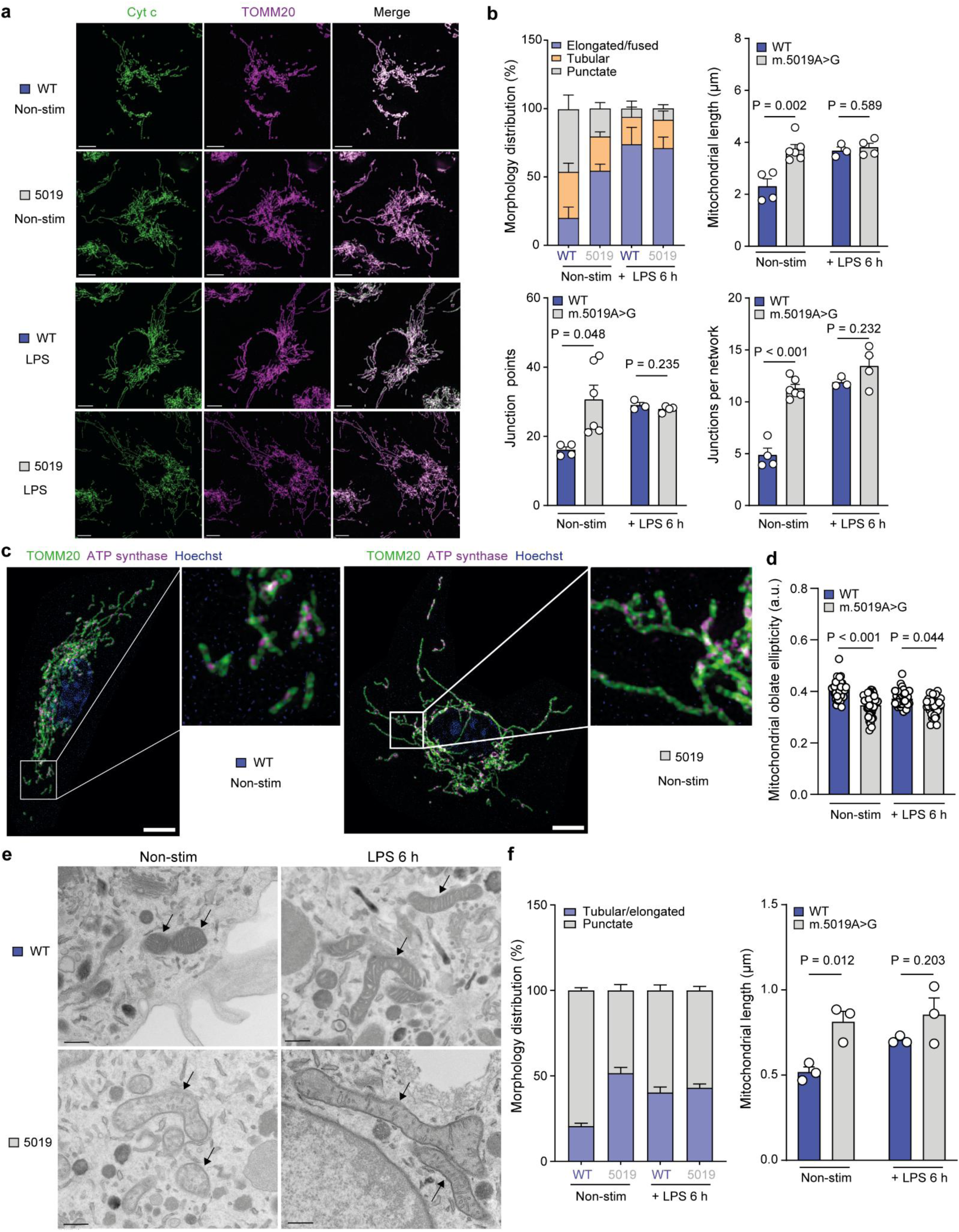
Increased mitochondrial elongation and disrupted cristae architecture in m.5019A>G primary macrophages Immunofluorescence staining of Cyt c and TOMM20 coupled to confocal microscopy (**a**) and morphology analysis (**b**) in non-stim (*n* = 4-6) and LPS-stimulated (*n* = 3-4; LPS 6 h) WT and m.5019A>G BMDMs. Scale bar 5 μm. Immunofluorescence staining of (**c**) TOMM20 and ATP synthase coupled to super-resolution microscopy and (**d**) morphology analysis of non-stim WT and m.5019A>G BMDMs (*n* = 3). Scale bar 5 μm. **e**, Transmission electron microscopy (TEM) and (**f**) morphology analysis of non-stim and LPS-stimulated WT and m.5019A>G BMDMs (*n* = 3; LPS 6 h). Scale bar 0.5 μm. Arrows indicate mitochondria. Representative images are shown. Data are mean ± s.e.m or ± s.d. *P* values calculated using two-tailed Student’s t-test for two group comparisons or one-way ANOVA corrected for multiple comparisons using the Kruskal-Wallis method.

### Combined respiratory chain deficiency in m.5019A>G macrophages

To identify the global molecular defects that arise as a consequence of the m.5019A>G mt-*Ta* mutation, we used unbiased data-independent acquisition (DIA) proteomic profiling (**Fig. 4a**) and transcriptomics (**Fig. 4d**). ORA of all differentially abundant proteins using gene ontology (GO) cellular compartment terms revealed a significant enrichment in mitochondrial proteins, including mitochondrial respiratory chain complexes and the mitoribosome (**Fig. 4b**). Further analysis revealed a significant decrease in the abundance of the nuclear-encoded structural subunits of CI, CIII, and CIV (**Fig. 4c**), likely due to a lack of core mtDNA-encoded protein subunits, consistent with quantitative proteomics performed on the cerebral cortex and liver of m.5019A>G mice^7^. No significant reduction was observed in CII or CV (**Extended data Fig. 4a**), in agreement with our respirometry and imaging analysis. In contrast, mitoribosome 28S and 39S subunits were increased in m.5019A>G macrophages likely as compensation for defects in mitochondrial translation (**Extended data Fig. 4b, c**). Gene set enrichment analysis (GSEA) following transcriptomic profiling, identified a decrease in OxPhos and basal IFN-I gene expression, and an increase in coagulation and hypoxia signatures in non-stim m.5019A>G macrophages (**Fig. 4d**). However, comparison of the Log_2_ fold change between the transcript and protein level of individual CI (**Fig. 4e**) and CIII and CIV (**Extended data Fig. 4d**) subunits revealed only a modest reduction on the transcriptional level when compared to protein abundance, indicating that the loss of nuclear-encoded OxPhos subunits occurs predominantly post-transcriptionally. This is consistent with reduced mitochondrial translation, which imbalances mito-nuclear OxPhos protein levels and lowers the assembly of fully functional respiratory chain complexes^37^. As such, impaired mitochondrial respiration and cristae disruption in m.5019A>G macrophages likely arise from a concurrent loss of mitochondrial and nuclear-encoded CI, CIII and CIV respiratory chain complexes.

**Figure 4.**
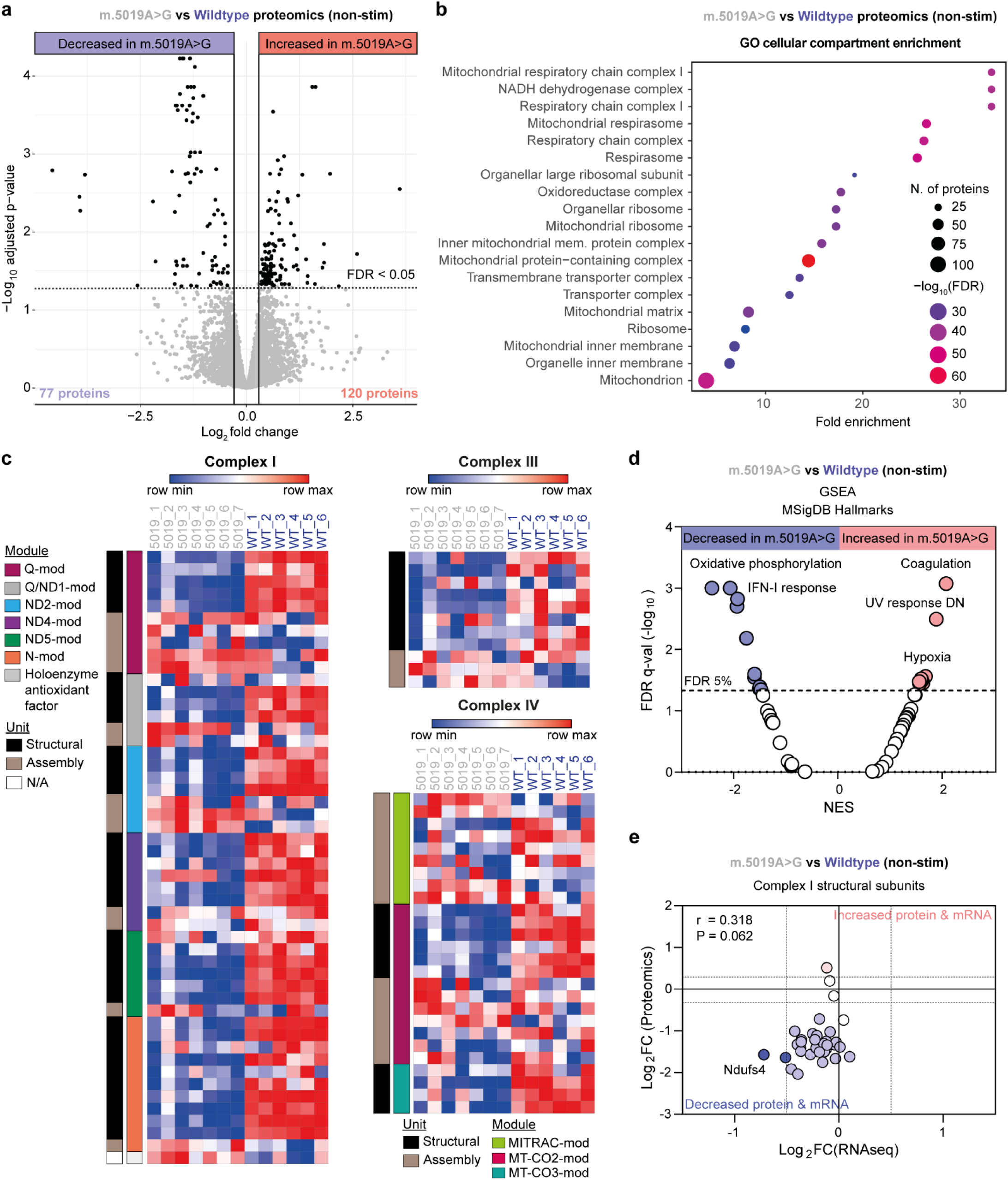
Decreased abundance of nuclear-encoded structural subunits of Complex I, III and IV in m.5019A>G primary macrophages **a,** Volcano plot of all differentially abundant proteins identified in non-stim m.5019A>G vs WT BMDMs (*n* = 6-7). **b**, Overrepresentation analysis (ORA) using gene ontology (GO) cellular compartment terms of all differentially abundant proteins identified in non-stim m.5019A>G vs WT BMDMs (*n* = 6-7). **c**, Heatmap of all identified Complex I (CI), CIII and CIV subunits and assembly factors in non-stim WT and m.5019A>G BMDMs (*n* = 6-7). **d**, Gene set enrichment analysis (GSEA) of RNA sequencing from non-stim WT and m.5019A>G BMDMs (*n* = 3). **e**, Comparison of Log_2_FC values of CI structural subunits from proteomics and RNA sequencing data with Pearson correlation and statistical analysis applied.

### TCA cycle remodelling and elevated NO production in m.5019A>G macrophages

The tricarboxylic acid (TCA) cycle undergoes functional remodelling during macrophage activation and is a key regulatory node for the synthesis of important immunomodulators, such as itaconate and NO^30,38–41^. Upon LPS stimulation, an inflammatory aspartate-argininosuccinate shunt (AAS) is induced in macrophages, driven by the increased expression of argininosuccinate synthetase 1 (ASS1) and the synthesis of argininosuccinate from aspartate^29,39^. This shunt is required to support the synthesis of arginine in the cytosol by argininosuccinate lyase (ASL), which is subsequently used by iNOS for NO production^29,39^. Analysis of TCA cycle metabolite abundance using LC-MS revealed a significant increase in α-ketoglutarate (α-KG), fumarate and malate levels in m.5019A>G versus WT macrophages (**Fig. 5a; left panel**). α-KG levels were further increased following LPS stimulation, which was accompanied by a significant reduction in succinyl-CoA levels, whereas itaconate levels remained similar between both genotypes (**Fig. 5a; left panel**). Synthesis of aspartate from oxaloacetate is dependent on mitochondrial respiration in proliferating cells^42,43^. However, a decrease in aspartate levels was only observed in m.5019A>G macrophages following LPS stimulation (**Fig. 5a; left panel**). Glutaminolysis is a major source of α-KG in macrophages^29,44^, and under short-term glutamine-depleted conditions, there was a more substantial reduction in TCA cycle metabolite levels in both non-stim and LPS-stimulated m.5019A>G macrophages (**Fig. 5a; right panel** and **extended data Fig. 5a**), suggesting an increased reliance on glutamine anaplerosis. In glutamine-depleted conditions, aspartate levels were reduced in both genotypes (**Fig. 5a; right panel** and **extended data Fig. 5a**), whilst a significant decrease in argininosuccinate was also observed in LPS-stimulated m.5019A>G macrophages. This shows that glutamine is an important nutrient for aspartate synthesis in macrophages, as previously reported^29^. Succinyl-CoA is synthesised from α-KG in the mitochondrial matrix by the oxoglutarate dehydrogenase complex (OGDHC)^38^. Comparisons of the α-KG/succinyl-CoA and α-KG/succinate ratios revealed a significant increase in both non-stim and LPS stimulated m.5019A>G macrophages compared to WT (**Fig. 5b** and **extended data Fig. 5b**) indicating reduced flux through the OGDHC. The OGDHC is composed of three subunits, E1 (OGDH), E2 (DLST) and E3 (DLD)^38^. Proteomic analysis revealed a significant and specific reduction in the E3 subunit in non-stim and LPS-stimulated m.5019A>G macrophages (**Fig. 5c**), consistent with increased α-KG and lower succinyl-CoA levels.

**Figure 5.**
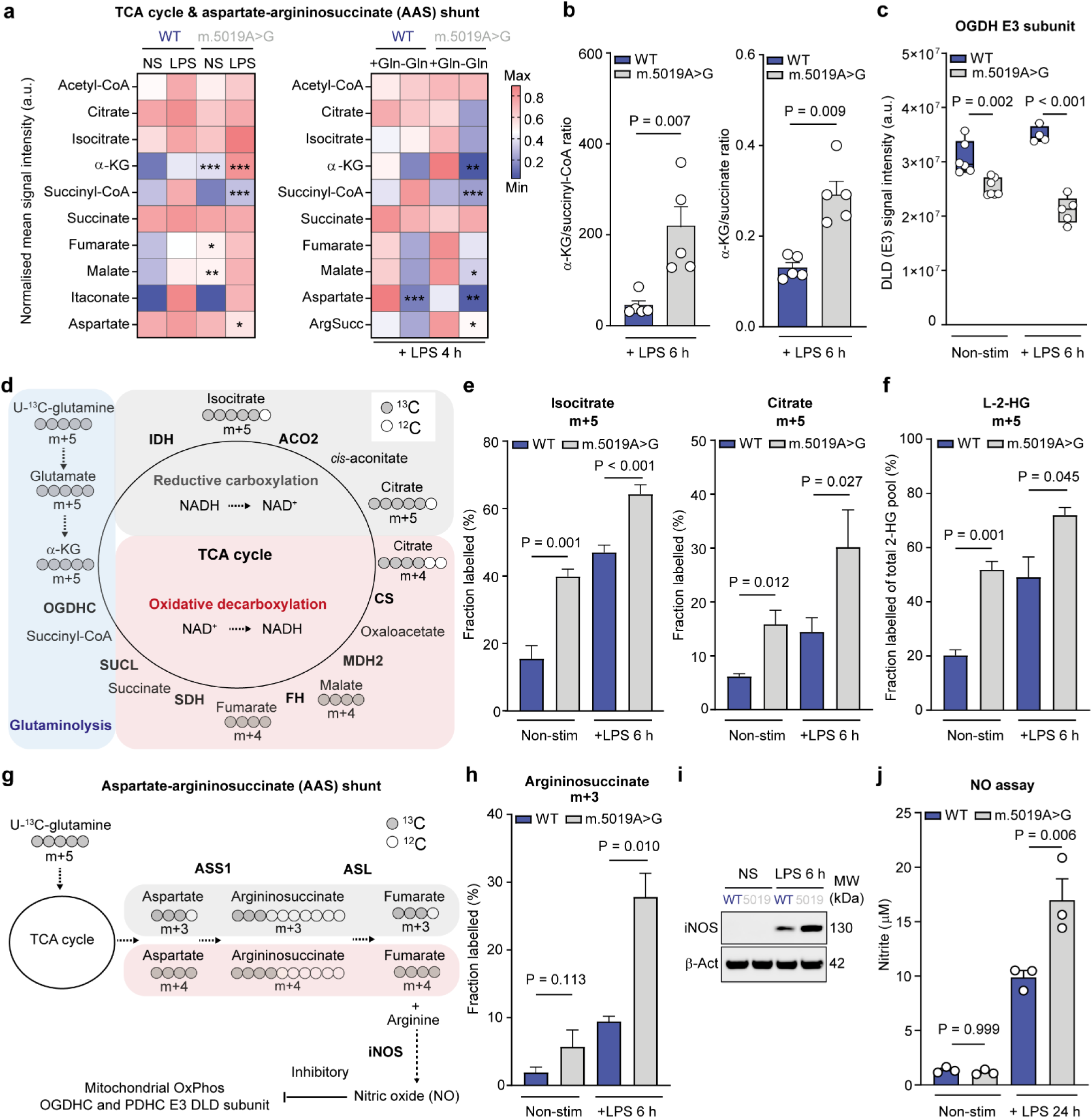
TCA cycle remodelling and increased nitric oxide production in m.5019A>G primary macrophages **a**, Heatmap comparing metabolite levels in non-stim and LPS-stimulated WT and m.5019A>G BMDMs (*n* = 5; LPS 6 h; left) and comparing metabolite levels in LPS-stimulated WT and m.5019A>G BMDMs in the presence or absence of glutamine (Gln) (*n* = 3; LPS 4 h; right). **b**, α-KG/succinyl-CoA and α-KG/succinate ratio in LPS-stimulated WT and m.5019A>G BMDMs (*n* = 5; LPS 6 h). **c**, OGDH E3 (DLD) subunit levels from proteomics in non-stim and LPS-stimulated WT and m.5019A>G BMDMs (*n* = 4-7). **d**, Schematic of U-^13^C-glutamine tracing into TCA cycle indicating oxidative versus reductive labelling patterns. **e**, m+5 labelling in isocitrate and citrate in non-stim and LPS-stimulated WT and m.5019A>G BMDMs (*n* = 5; LPS 6 h). **f**, m+5 labelling in L-2-hydroxyglutarate (L-2-HG) levels in in non-stim and LPS-stimulated WT and m.5019A>G BMDMs (*n* = 3; LPS 6 h). **g**, Schematic of U-^13^C-glutamine tracing into aspartate-argininosuccinate shunt (AAS) indicating oxidative versus reductive labelling patterns and NO production. **h**, m+3 labelling in argininosuccinate in non-stim and LPS-stimulated WT and m.5019A>G BMDMs (*n* = 5; LPS 6). **i**, iNOS protein levels (*n* = 5; LPS 6 h; representative blot shown) and (**j**) nitrite levels (*n* = 3; LPS 24 h) in cell culture medium (CCM) in non-stim and LPS-stimulated WT and m.5019A>G BMDMs. Data are mean ± s.e.m. *P* values calculated using two-tailed Student’s t-test for two group comparisons or multiple unpaired t-tests corrected for multiple comparisons using Holm-Sidak method. **\*\*\*** *P* < 0.001 **\*\*** *P* < 0.01 **\*** *P* < 0.05.

The TCA cycle can operate in the forward (oxidative) and reverse (reductive) direction^10,45^. Oxidative metabolism is important for NADH generation, whereas reductive metabolism is associated with NADH consumption and macromolecule biosynthesis in respiratory-deficient cancer cells^46,47^. Impairments in CI-dependent respiration and reduced OGDHC levels are known to increase reductive carboxylation in proliferating cells^48^. Typically, hypoxic and respiratory-deficient cancer cells engage reductive carboxylation due to a decrease in the NAD^+^/NADH ratio and associated reductive stress^46,47^. However, no significant differences in the whole cell NAD^+^/NADH ratio were observed between WT and m.5019A>G macrophages (**Extended data Fig. 5c**). We subsequently performed stable isotope-assisted U-^13^C-glutamine tracing coupled to LC-MS, which can assess distinct oxidative and reductive metabolism to reveal compartmentalised redox states (**Fig. 5d** and **extended data Fig. 5d-f**). Comparison of the m+5/m+3 ratio of α-KG confirmed reduced oxidative TCA cycle activity in m.5019A>G macrophages (**Extended data Fig. 5d**). Further analysis revealed a significant increase in m+5 labelling and reduced m+4 labelling in isocitrate and citrate in non-stim m.5019A>G macrophages (**Fig. 5e** and **Extended data Fig. 5e**), which was further increased following LPS-stimulation. This data confirms the presence of increased mitochondrial reductive stress and reduced flux through OGDHC in m.5019A>G macrophages. When compared to non-stim conditions, LPS stimulation also increased reductive m+5 labelling in isocitrate and citrate in WT macrophages in agreement with a previous study^49^. 2-hydroxyglutarate (2-HG) is a metabolite synthesised from α-KG under conditions of OGDHC deficiency and reductive stress^48^. Consistent with this, 2-HG levels (**Extended data Fig. 5g**) and m+5 labelling from U-^13^C-glutamine (**Extended data Fig. 5h**) were also increased. 2-HG exists as two enantiomers, L-2-HG and D-2-HG, which can only be differentiated by LC-MS after derivitisation^48^. Indeed, we observed a significant increase in L-2-HG m+5 labelling in m.5019A>G macrophages (**Fig. 5f** and **extended data Fig. 5h**), consistent with reports of OGDHC deficiency^48^. U-^13^C-glutamine tracing also revealed a significant increase in m+3 labelled argininosuccinate (**Fig. 5g, h**), fumarate and malate (**Extended data Fig. 5f**) in LPS-stimulated m.5019A>G macrophages, which are derived from the reductive synthesis of aspartate (m+3) (**Fig. 5g** and **extended data Fig. 5f**). Therefore, reductive carboxylation is a feature of not only hypoxic and respiratory-deficient cancer cells, but also respiratory-deficient macrophages, where it fuels the inflammatory AAS. Finally, LPS-induced iNOS levels (**Fig. 5i**) and NO production (**Fig. 5j**) were higher in m.5019A>G macrophages, as measured by Western blot and nitrite concentration in the CCM, respectively. This data is consistent with reports that IFN-β accelerates the induction of iNOS by LPS^50^. NO is an established inhibitor of mitochondrial respiratory chain complexes, the pyruvate dehydrogenase complex (PDHC) and OGDHC, which act to restrict pro-inflammatory IL-1β and IL-6 production^38,51,52^ and promote IFN-β release^41^ in macrophages. In addition, elevated α-KG levels have previously been reported to weaken pro-inflammatory cytokine responses and contribute to endotoxin tolerance^44^. As such, this combination of TCA cycle remodelling and elevated NO production is likely a contributing factor to the observed imbalance in m.5019A>G macrophage cytokine responses.

### Glycolytic reprogramming in m.5019A>G macrophages

Interestingly, no changes in whole cell ATP/AMP or ATP/ADP ratios were observed in non-stim m.5019A>G macrophages (**Extended data Fig. 6a**), despite impaired mitochondrial respiration and oxidative TCA cycle metabolism. This suggested that these macrophages could compensate for the bioenergetic defects. To understand this, we turned to our transcriptomic and proteomics data and identified a significant increase in signatures for HIF-1α signalling, hypoxia, glycolysis and glutathione metabolism (**Fig. 4d and Fig. 6a-c**). Macrophages engage in aerobic glycolysis following inflammatory activation^38,51,53^, which is thought to support cellular ATP synthesis, lactate production by lactate dehydrogenase (LDH), and NAD^+^ regeneration in the face of impaired mitochondrial respiration. Phenotyping of WT and m.5019A>G macrophages using an extracellular flux analyser highlighted a reduction in OCR and concomitant increase in the extracellular acidification rate (ECAR), under non-stim and LPS-stimulated conditions in m.5019A>G macrophages (**Fig. 6d**). A glycostress test confirmed the increase of glycolysis in non-stim m.5019A>G macrophages (**Fig. 6e**). Inhibition of CV with oligomycin increased max glycolytic capacity in WT macrophages but failed to do so in m.5019A>G macrophages, causally linking impaired mitochondrial respiration to increased aerobic glycolysis (**Fig. 6e**). Intracellular lactate measurements in glucose replete-and depleted-conditions (**Fig. 6f**) and extracellular lactate measurements (**Fig. 6g**) confirmed increased aerobic glycolysis in non-stim m.5019A>G macrophages, but extracellular lactate levels did not increase further following LPS stimulation suggesting they were already working at max glycolytic capacity at this timepoint. Glucose deprivation also had opposing impacts on TCA cycle metabolite abundance in m.5019A>G macrophages compared to WT macrophages (**Extended data Fig. 6b**), which demonstrates a reduced capacity of respiratory-deficient mitochondria in m.5019A>G macrophages to adapt to glucose restriction. Finally, in support of the proteomics data, increased total glutathione levels were also observed in m.5019A>G macrophages (**Fig. 6h**), consistent with previous reports of mitochondrial respiratory disruption^29,54^. Increased aerobic glycolysis and lactate synthesis in m.5019A>G macrophages prior to inflammatory activation likely supports a recently identified role for lactate in the immunosuppression of pro-inflammatory cytokines, including *Il1b* and *Il6*^55^. In addition, the combination of reduced mitochondrial respiration and increased aerobic glycolysis is reminiscent of that reported in LPS-tolerant macrophages, whereby mitohormetic adaptations restrain pro-inflammatory macrophage responses^24,56^. Overall, our data suggests that the combined impact of the m.5019A>G mutation on mitochondrial function and cellular metabolic reprogramming reinforces a metabolic state in macrophages that favours IFN-I production.

**Figure 6.**
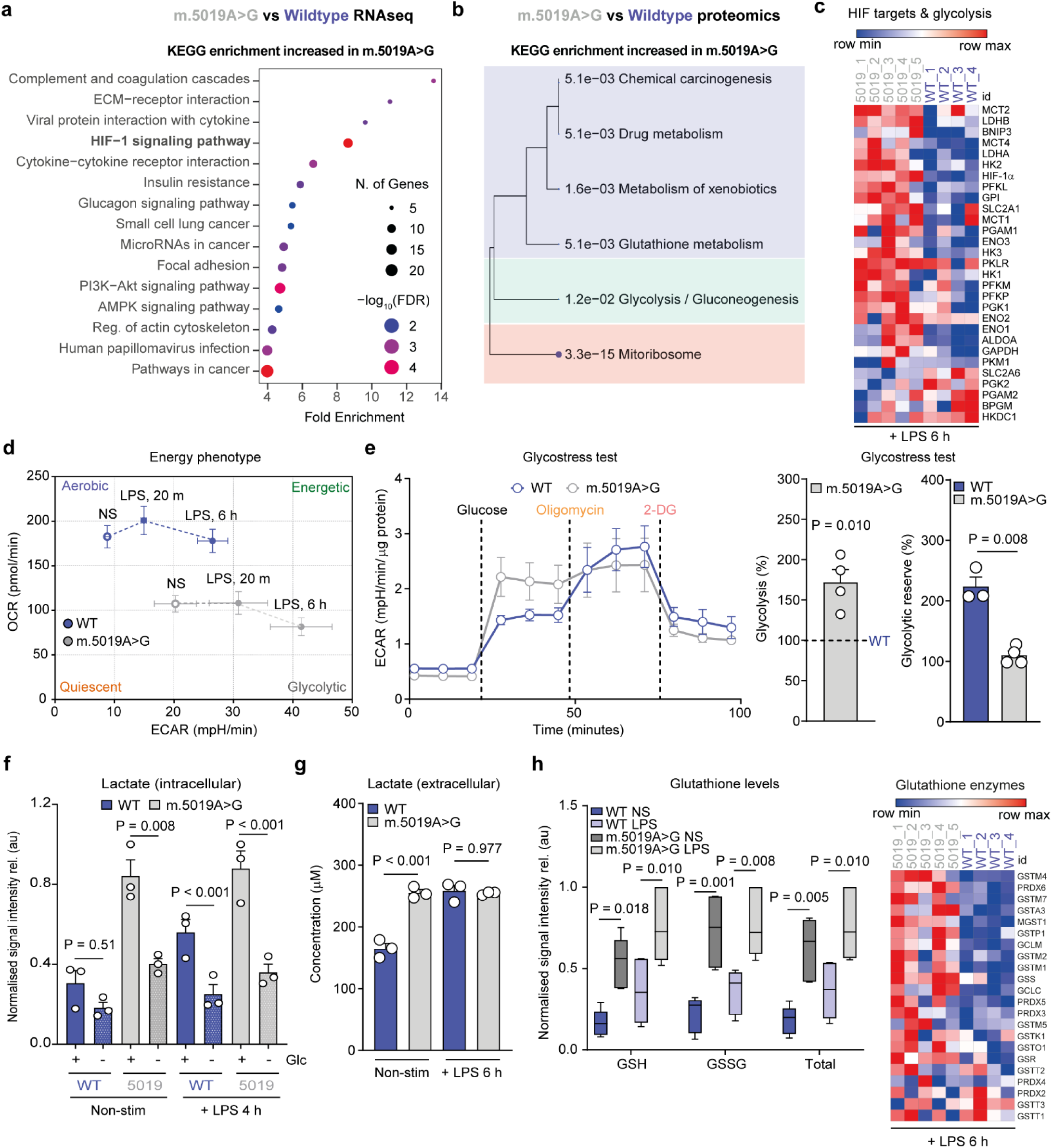
Increased aerobic glycolysis and glutathione in m.5019A>G primary macrophages ORA using KEGG terms of (**a**) all differentially expressed genes from RNA sequencing (*n* = 3) and (**b**) differentially abundant proteins (*n* = 6-7) increased in non-stim m.5019A>G vs WT BMDMs. **c**, Heatmap of HIF-1α targets and glycolytic enzymes from proteomics in LPS-stimulated WT and m.5019A>G BMDMs (*n* = 4-5; LPS 6 h). **d**, Oxygen consumption rate (OCR) and extracellular acidification rate (ECAR) measurements in non-stim and LPS-stimulated WT and m.5019A>G BMDMs (*n* = 6-7). **e**, ECAR measurements using the Glycostress test in non-stim WT and m.5019A>G BMDMs (*n* = 3-4). **f**, Intracellular lactate measurements in non-stim and LPS-stimulated WT and m.5019A>G BMDMs in the presence or absence of glucose (Glc) (*n* = 3; LPS 4 h). **g**, Extracellular lactate measurements in non-stim and LPS-stimulated WT and m.5019A>G BMDMs (*n* = 3; LPS 6 h). **h**, Glutathione metabolite levels (non-stim and LPS-stimulated) (*n* = 5) and enzyme abundance (*n* = 4-5; LPS 6 h) WT and m.5019A>G BMDMs. Data are mean ± s.e.m. *P* values calculated using two-tailed Student’s t-test for two group comparisons or one-way ANOVA corrected for multiple comparisons using Tukey method.

### Elevated type I IFN levels in m.5019A>G mice

Finally, to determine whether the innate immune response of m.5019A>G mice was distinct from WT mice, we injected a sub-lethal dose of LPS intraperitoneally to collect serum for cytokine and chemokine analysis and monitor sickness behaviour (**Fig. 7a**). Consistent with elevated IFN-β in m.5019A>G macrophages *ex vivo*, we observed a robust and significant increase in serum IFN-β levels in female and male m.5019A>G mice (**Fig. 7b, c**). Olink profiling also revealed a significant increase in IFN-α2 levels in both female and male m.5019A>G mice (**Fig. 7d, e**), while TNF-α and IL-10 were also significantly higher in female m.5019A>G mice. Sickness behaviour was also noticeably more severe in m.5019A>G mice (**Fig. 7f, g)**. Importantly, this data confirms that a heteroplasmic mtDNA mutation can perturb systemic innate immune responses *in vivo*.

**Figure 7.**
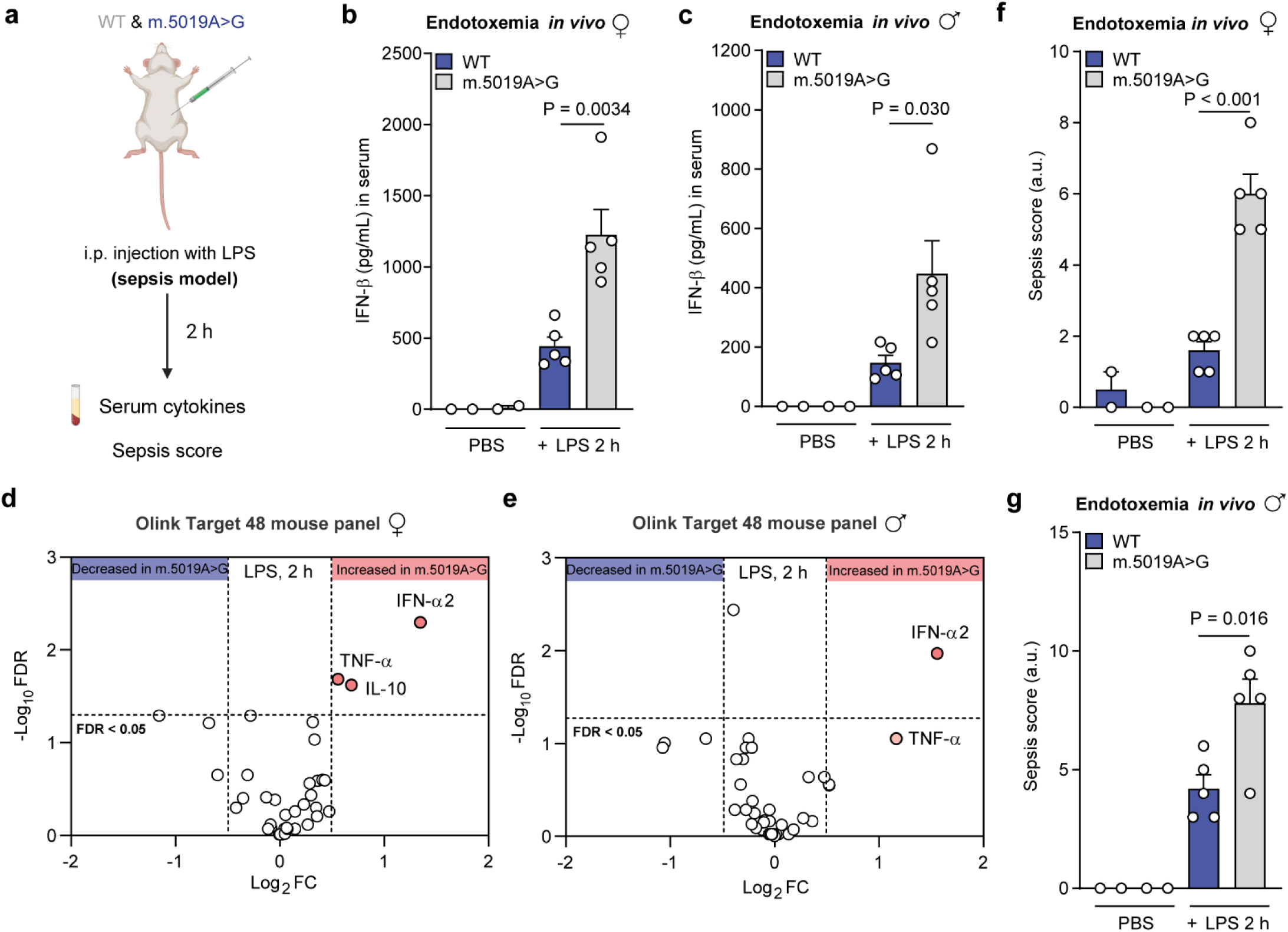
Elevated type I IFN levels in m.5019A>G mice following an *in vivo* LPS challenge **a,** Schematic of *in vivo* LPS endotoxemia model experiment. IFN-β levels in serum of (**b**) female and (**c**) male m.5019A>G mice injected intraperitoneally (i.p.) with PBS (*n* = 2; 2 h) or LPS (*n* = 5; 2.5 mg/kg; 2 h). Olink target T48 mouse cytokine and chemokine profiling of serum in LPS-stimulated (**d**) female and (**e**) male m.5019A>G mice vs WT mice (*n* = 5; LPS 2 h). Sepsis score in (**f**) female and (**g**) male m.5019A>G and WT mice injected intraperitoneally (i.p.) with PBS (n = 2; 2 h) or LPS (n = 5; 2.5 mg/kg; 2 h). Data are mean ± s.e.m. *P* values calculated using two-tailed Student’s t-test for two group comparisons or multiple unpaired t-tests corrected for multiple comparisons two-stage step-up method of Benjamini, Krieger and Yekutieli method.

## Discussion

An increasing number of clinical reports have identified that patients with primary mitochondrial disease are more susceptible to severe infections and sepsis^19–21,57^. In addition, previous pre-clinical studies have established that deletion or mutation of nuclear-encoded mitochondrial proteins can lead to premature death, sensitivity to endotoxemia, accelerated ageing, and hyperinflammatory macrophage responses^1–5,28,58^. A key focus of our study was to understand whether any perturbations in innate immune signalling were evident in macrophages or mice with inherited heteroplasmic mtDNA mutations, which constitute the largest proportion of primary mitochondrial disease cases^13^. To our knowledge, the only other report of the impact of a heteroplasmic mtDNA mutation on murine macrophages used the m.5024C>T mt-*Ta* mouse model^59^. In line with our findings, m.5024C>T macrophages could tolerate high mutational burdens with no evidence of purifying selection during differentiation despite reduced mitochondrial OxPhos^59^. MHC class II levels were also decreased in m.5024C>T macrophages, consistent with our observations in m.5019A>G macrophages. However, this important study predominantly focused on purifying selection against the m.5024C>T mutation in antigen-stimulated T and B cells of the adaptive immune system and did not address our core research question.

The key finding of our study is that IFN-I release is elevated in m.5019A>G macrophages *ex vivo* and m.5019A>G mice *in vivo* following an LPS challenge, consistent with reports in *Polg*^D257A^ mutator mice^1,3^. This provides compelling evidence to support the idea that mitochondrial dysfunction arising from inherited mtDNA mutations perturb innate immune signalling. IFN-I are well known for their important roles in anti-viral and anti-bacterial host defence. However, paradoxically, they are also a primary driver of tissue damage and pathology during microbial infection or in autoimmune disorders, often via host immunosuppression and antagonism of pro-inflammatory cytokine responses^1,60,61^. This growing body of literature clearly demonstrates that IFN-I production needs to be tightly controlled to mitigate detrimental impacts on physiology^32^. In support of this, increased IFN-I signalling is the defining feature of a set of Mendelian inborn errors of immunity termed type I interferonopathies first described in 2011^62^. Increased IFN-I signatures have now also been observed in primary mitochondrial disease patients and may represent a novel clinical biomarker^17,63^. In addition, primary mitochondrial disease patients are more susceptible to vaccine preventable infections, while sepsis and SIRS are a major cause of morbidity and mortality^8,19,20,57^. In some clinical reports, intercurrent infection is also reported to precipitate the onset of neurological symptoms, a key feature of many primary mitochondrial disease subtypes caused by inherited mtDNA mutations^21^. Given that dysregulated IFN-I signalling has been identified as both a driver of sepsis^32,60^ and neurological disease in type I interferonopathies^64^, this may underlie the susceptibility of mitochondrial disease patients to severe and recurrent infection, in addition to other clinical disease manifestations. Overall, this growing body of evidence suggests that inherited mtDNA mutations may give rise to an inborn error of immunity, thus impacting the response of mtDNA disease patients to both environmental pathogen exposure and maintenance of tissue homeostasis, with important clinical implications for disease management and treatment.

Mechanistically, elevated IFN-I responses downstream of mitochondrial dysfunction have previously been linked to mtDNA release, while other studies have shown that cytosolic mtRNA release can also enhance IFN-β production in respiratory-deficient macrophages following LPS stimulation^29,30^. Indeed, we observe disrupted mitochondrial cristae architecture and increased cytosolic mtDNA and mtRNA levels in m.5019A>G macrophages, which may explain this imbalance toward IFN-I following LPS stimulation. However, it is important to note that both cytosolic mtDNA and mtRNA levels were higher in non-stimulated m.5019A>G macrophages with no apparent increase in macrophage activation. In fact, basal IFN-I gene expression was modestly reduced in macrophages in the resting state and no appreciable levels of IFN-β were detected in the serum of uninfected m.5019A>G mice. A wide variety of intracellular bacterial and viral pathogens promote the release of mtDNA to fully engage the host innate immune response following infection^9^. Our data suggests that these mitochondrial signals, by themselves, are insufficient to drive IFN-I expression in macrophages in the absence of an infection stimulus in this context. This may be explained by increased intracellular lactate levels, which have been reported to restrict RIG-I-like receptor (RLR) and cGAS-mediated sensing of dsRNA and DNA, respectively^65,66^. Alternatively, an as yet ill-defined threshold may need to be bypassed to promote inflammation in macrophages in the absence of infection. IFN-β is reported to accelerate the induction of iNOS by LPS^50^, while elevated NO production is an established inhibitor of respiratory chain complexes, PDHC and OGDHC in inflammatory macrophages, which acts to impair mitochondrial respiration and promote lactate synthesis^38,41,51,52^. The outcome of this combined metabolic state serves to restrict pro-inflammatory IL-1β and IL-6 production^38,51,52,56^ and promote IFN-β release^41^. This may explain the higher levels of IFN-β after prolonged LPS stimulation in m.5019A>G macrophages. As such, the observed imbalance in the m.5019A>G macrophage cytokine response likely arises from a combination of mtDNA mutation-driven impairment in mitochondrial respiration and toll-like receptor 4 (TLR4)-driven metabolic reprogramming. In conclusion, this heteroplasmic mtDNA mutation model provides an excellent opportunity to explore mechanistic links between inherited mtDNA mutations, IFN-I signalling, and disease pathology in future studies, with the goal of identifying therapeutic strategies for clinical intervention.

## Materials and methods

### Animals

All mouse experiments and breeding were carried out in accordance with the UK Animals (Scientific Procedures) Act, 1986 (Home Office PPL no. PP1740969) and EU Directive 2010/63/EU. All experiments followed ARRIVE guidelines and procedures were approved by the University of Cambridge Animal Welfare and Ethical Review Body (AWERB) Committee. Wildtype (WT) mice were purchased from Charles River Laboratories, UK. The m.5019A>G mouse strain (Allele symbol: mt-T^am2Jbst^, MGI ID: 6860509) was generously provided from the colony of Patrick F. Chinnery. The mice were provided through an MTA with James. B Stewart. All mice used in this study were on the C57BL/6J background. WT and m.5019A>G mice were age and sex matched for all experiments. Mice were kept in individually-ventilated cages (Tecniplast) at 20-24°C, 45-65% humidity, with a 12 h light-dark cycle and with ad libitum access to SAFE 105 universal diet (Safe Diets) and water.

### Generation of murine bone marrow-derived macrophages (BMDMs)

Mice aged 8-33 weeks old were euthanised by cervical dislocation and death was confirmed by exsanguination. Bone marrow was harvested from the tibia and fibula. Cells were pelleted by centrifugation at 1500 rpm for 5 mins and red blood cells were lysed using a hypotonic red blood cell lysis buffer (Abcam, ab204733). Cells were pelleted by centrifugation at 1500 rpm for 5 min and filtered through a 70 µm nylon mesh. Obtained cells were differentiated in DMEM (Gibco^TM^) containing L-929 (ATCC^®^) conditioned medium (20%), Foetal Bovine Serum (10%) (FBS, Gibco™ A5670701) and penicillin-streptomycin (1%) (Gibco™, 15070-063) for 6 days. Media was changed on day 4. On day 6, BMDMs were scraped and counted using trypan blue, and then plated in DMEM with L929-conditioned medium (10%), FBS (10%) and penicillin-streptomycin (1%). BMDMs were plated at 0.5 x 10^6^ cells/well (1 mL total volume) in 12-well cell culture plates (150628, ThermoFisher Scientific) and left overnight to adhere at 37°C in a de-humidified incubator (21% O_2_, 5% CO_2_), unless otherwise stated. BMDMs were activated with LPS from *Escherichia coli*, serotype EH100 (Alexis) used at a final concentration of 100 ng/mL.

### Flow cytometry analysis

BMDMs were plated at 1 x 10^6^ cells/well in 6-well cell culture plates (2 mL total volume) and left to adhere overnight at 37°C in a de-humidified incubator (21% O_2_, 5% CO_2_). For macrophage differentiation, F4/80 cell surface marker (Invitrogen, 12-4801-80) was used according to manufacturer’s instruction. For mitochondrial mass and membrane potential measurements, cells were incubated with MitoTracker Green FM (M7514, ThermoFisher Scientific) or Tetramethyl rhodamine Methyl Ester Perchlorate (TMRM, 11550796, Invitrogen) according to the manufacturer’s instructions. Cells were then washed twice with PBS, scraped and resuspended in DMEM with FBS (1%). Flow cytometry analysis was performed on a BD Fortessa flow cytometer and further analysed using FlowJo software.

### mtDNA heteroplasmy measurements

At weaning, ear skin biopsies were taken from all heteroplasmic m.5019A>G mice to allow each animal to be assigned a reference heteroplasmy value. DNA was extracted from ear skin biopsy samples, bone marrow and BMDMs using the DNeasy Blood & Tissue kit (69504, Qiagen) according to the manufacturer’s protocol and quantified using a NanoDrop spectrophotometer (ThermoFisher Scientific). Heteroplasmy measurements were determined by pyrosequencing using the Q48 Autoprep or Q24 vacuum workstation systems (Qiagen) as previously described^7^. In brief, a section of the mitochondrial genome containing the mutation of interest was amplified from approximately 25 ng of DNA using the Pyromark® PCR kit (978703, Qiagen) and primers (Integrated DNA technologies (IDT)) designed according to the manufacturer’s instructions. Following PCR, 3 µL of Pyromark® magnetic beads (974203, Qiagen) were loaded with 10µL of biotinylated PCR product onto Pyromark® Q48 disks (974901, Qiagen). Disks were run on a Q48 Pyromark® sequencer (Qiagen) using an allele quantification assay using the following dispensation order: TG/AAGGAC/TTGTAAG using a sequencing primer. Heteroplasmy was then called using Pyromark® analysis software (Qiagen) and reported as the percentage of mutant base present in the sample.

### Mitochondrial DNA copy number and levels measurement

Mitochondrial DNA (mtDNA) copy number was quantified by digital droplet PCR (ddPCR) and levels by quantitative PCR (qPCR). DNA was isolated using DNeasy Blood & Tissue Kit (69504, Qiagen) according to the manufacturer’s instructions. The DNA concentration of each sample was measured using a Nanodrop ND-8000 UV-visible spectrophotometer (ThermoFisher) and then adjusted to 10 ng/μL.

For ddPCR, a mixture of 1μL DNA and 21μL reaction mixture was made up and placed in a 96-well ddPCR plate (Bio-Rad). The plate was then sealed, vortexed for 20 secs, centrifuged with a pulse spin and placed on a chilled block (4°C) (Bio-Rad). Droplets were generated using a QX200 AutoDG droplet generator with corresponding automated droplet-generating oil (1864110, Bio-Rad), DG32 automated droplet generator cartridges (1864108, Bio-Rad) and pipette tips for AutoDG system (1864120, Bio-Rad). Each item was placed in the appropriate place in the instrument, the plate was unsealed and placed on the chilled block, in the droplet generator. Upon completion, the plate was removed and sealed with foil (1814040, Bio-Rad) using a plate sealer (Bio-Rad) set at 180°C for 5 secs. The plate was stored on ice until the next step. ddPCR was run on a C1000 Touch Thermal Cycler (BioRad) programmed to 95°C for 30 secs and 60°C for 1 min repeated for 40 cycles followed by 90°C for 10 mins. Temperature change ramp rates were set at 2°C/sec. The plate was stored at 4°C until analysis. Droplets were read by a QX200 Droplet Reader (Bio-Rad) using the following experimental settings: absolute quantification, rare event detection, copy number variation and supermix for probes with no dUTP. The results were analysed using QuantaSoft analysis software (Bio-Rad) and the average mtDNA copy number (HEX probe) was normalised to the nuclear DNA copy number (FAM probe). Primers and probe used below:

For qPCR, primers specific for the mitochondrial *D-loop* region and a specific region of mtDNA that is not inserted into nuclear DNA (*non-NUMT*) were used, as previously described^67,68^. *Tert* and *B2m* primers were used for normalisation. The reaction was performed in a MicroAmp optical 96-well reaction plate (N8010560, Applied Biosystems) using a 96-well QuantStudio^TM^ 3 PCR machine and PowerUp™ SYBR™ Green Master Mix (A25741, Applied Biosystems). Data for normalised *D-loop* and *non-NUMT* were combined to assess mtDNA levels. Primer sequences are as follows:

### Extracellular flux analysis (Seahorse)

BMDMs were seeded at 0.2 x 10^6^ cells/well in a Seahorse XFe24 well plate and left to adhere overnight at 37°C in a de-humidified incubator (21% O_2_, 5% CO_2_). Oxygen consumption rate (OCR) and Extracellular acidification rate (ECAR) were analysed using an Energy Phenotype test, Seahorse XF Mito Stress test (103015-100, Agilent) or Seahorse XF Glycolysis Stress test (103020-100, Agilent) according to the manufacturer’s instructions. In brief, a utility plate containing XF Calibrant fluid (100840-000, Agilent), together with the sensor cartridge, was placed in a CO_2_-free incubator at 37°C overnight. The following day, the medium was replaced with 500 μL of XF DMEM medium pH 7.4 (103575-100, Agilent) supplemented with glucose (10 mM), glutamine (2 mM), and sodium pyruvate (1 mM) for the Energy Phenotype and Seahorse XF Mito Stress test or glutamine (2 mM) for the Seahorse XF Glycolysis Stress test. The cell culture plate was then placed in a CO_2_-free incubator at 37°C for 45-60 mins prior to analysis on Seahorse XFe24 Analyser (Agilent). Seahorse XF Mito Stress Test Kit inhibitors (2 μM Oligomycin A, 1.0 μM FCCP, 0.5 μM Rotenone and 0.5 μM Antimycin A, final well concentration), Seahorse XF Glycolysis Stress Test Kit (10 mM glucose, 2 μM Oligomycin A and 50 mM 2-deoxyglucose (2-DG), final well concentration) and Energy Phenotype (200 ng/mL lipopolysaccharide (LPS), final well concentration) were added to the appropriate ports of the injector plate. After calibration of the utility plate and sensor cartridge, the cell culture plate was analysed on the Seahorse XF24 analyser using default test settings. All results were acquired with Wave software (Agilent) and analysed with Seahorse XF test report generators according to manufacturer’s instructions. For Seahorse XF Mito Stress test and XF Glycolysis Stress test, protein concentration was determined at experimental endpoints for normalisation of OCR and ECAR data to total protein (μg) using a Pierce™ BCA Protein Assay Kit (23225, ThermoFisher Scientific).

### Respirometry analysis (Oroboros)

Oxygen consumption rate (OCR) in BMDMs was measured using an Oxygraph-2k high-resolution respirometer (Oroboros Instrument, Innsbruck, Austria) in 2 ml glass chambers at a constant temperature of 37°C and stirrer speed of 750 rpm. Oxygen flux (*J*O_2_), which is directly proportional to OCR, was continuously recorded with a 2 sec sampling rate using DatLab software 6.1 (Oroboros Instruments, Austria). Calibration at air saturation was carried out every day prior to experimentation, and all data were corrected for background instrumental *J*O_2_ in accordance with the manufacturer’s instructions. BMDMs were counted directly before resuspending in mitochondrial respiration medium (MIRO5) at 1 x 10^6^ cells/mL. WT and m.5019A>G BMDMs were then added to individual Oxygraph-2k chambers to enable paired comparisons and allowed to reach a stable baseline OCR prior to stimulations. All reagents injected into the chambers (Digitonin, Malate, Glutamate, ADP, Cytochrome c, Succinate, FCCP and Rotenone) were warmed to room temperature prior to addition. Digitonin (14952-100mg-CAY, Cayman Chemical) was used to permeabilise the plasma membrane. Malate (M1000, Sigma) and glutamate (G1626, Sigma) were injected to determine complex I (CI) leak activity. ADP (A2754, Sigma) was subsequently injected to determine CI-dependent OxPhos. Cytochrome c (C7752, Sigma) was injected to ensure integrity of the mitochondrial membrane. Succinate (S2378, Sigma) was injected to determine maximum OxPhos. FCCP (15218, Cambridge Bioscience) was injected to dissipate membrane potential to assess maximum electron transfer capacity (ETC). Rotenone (HY-B1756, Cambridge Bioscience) was subsequently injected to inhibit CI activity to determine CII ETC. The flux control ratio for CI was assessed from CI OxPhos and maximum OxPhos, while for CII was assessed from CII ETC and maximum ETC.

### ^35^S-methionine labelling of mitochondrial translation products

BMDMs were plated at 0.5 x 10^6^ cells/mL (2 mL total volume) in a 6-well dish and left to adhere overnight at 37°C in a de-humidified incubator (21% O_2_, 5% CO_2_). In order to label newly synthesised mitochondrial proteins, the previously published protocol was used^69^. Briefly, cells were incubated in methionine/cysteine-free medium for 10 mins before being replaced with methionine/cysteine-free medium containing 10% dialysed FCS and emetine dihydrochloride (100 μg/ml) to inhibit cytosolic translation. Following a 20 min incubation, 120 μCi/ml of [^35^S]-methionine was added, and the cells were incubated for 30 min. After washing with PBS, cells were lysed, and 30 μg of protein was loaded on 10–20% Tris-glycine SDS-PAGE gels. Coomassie staining for total protein was also performed. Dried gels were visualized with a PhosphorImager system. The ^35^S-methionine signal intensity was subsequently normalised to the Coomassie staining to assess mitochondrial translation.

### Confocal microscopy

BMDMs were plated at 0.25 x 10^6^ cells/well (0.5 mL total volume) on coverslips in a 24-well plate and left to adhere overnight at 37°C in a de-humidified incubator (21% O_2_, 5% CO_2_). Cells were stimulated as indicated. Cells were subsequently washed three times with PBS and fixed with paraformaldehyde (PFA) (4%) for 15 mins at room temperature. After washing, cells were permeabilised and non-specific binding was blocked with 0.1% Triton X-100, 3% BSA in PBS for 30 mins. Incubation with primary antibodies for TOMM20 (11802-1AP, Proteintech) and Cytochrome c (556432, BD Biosciences) was performed overnight at 4°C in blocking buffer. Incubation with Alexa Fluor 488 or 561 secondary antibodies (ThermoFisher Scientific) was performed for 30 mins at room temperature. The slides were mounted using ProLong Gold Antifade Mountant (P36982, Invitrogen). All images were acquired using a Zyla 4.2 PLUS sCMOS camera attached to an Andor DragonFly 500 confocal spinning disk mounted on a Nikon Eclipse TiE microscope using a CFI Plan Apochromat lambda 100X oil immersion objective and using the Fusion user interface (Andor). Images were prepared and analysed with Fiji ImageJ (NIH). Mitochondrial morphology percentage per cell were done as previously described^70^. Mitochondrial length quantifications were done using a published Mito-Morphology macro^71^ and a Fiji ImageJ macro script to determine mitochondrial objects, mitochondrial networks, junction points and junctions per network using preselected regions of interest (ROIs). At least 20 cells per mouse per condition were analysed in each experiment.

### Super-resolution microscopy

BMDMs were plated at 0.5 x 10^6^ cells/mL (0.5 mL total volume) on coverslips in a 24-well plate and left to adhere overnight at 37°C in a de-humidified incubator (21% O_2_, 5% CO_2_). Following indicated treatments, BMDMs were fixed in PFA (5%), pH 7.4 for 15 min at 37°C and then washed three times with PBS. After blocking with FBS (10%) for 30 mins, macrophages were incubated with ATP Synthase (MAB3494, Merck) and TOMM20 (ab232589, Abcam) primary antibodies at 1:500 in FBS (5%) overnight at 4°C. Cells were then washed with FBS (5%) three times and incubated with fluorescent secondary antibodies at 1:1000 for 1 h at room temperature with shaking, protected from light, and then washed three times with PBS. Cells were then incubated with Hoechst 33342 (1:500 in PBS) for 1 h at room temperature, washed three times with PBS, mounted onto glass slides using mounting medium (ProLong Diamond), left to dry for 12 h at room temperature, and then stored at 4 °C until imaging. Fixed cell super-resolution images for analysis of mitochondrial morphology in 3D were obtained with the Zeiss Elyra7 lattice SIM, using the Plan-Apochromat 63x/1.4 Oil DIC M27 objective with 15 phases and 0.091 μm intervals. Images were acquired with 20 ms exposure time with 405 nm (20.0%), 488 nm (4.0%), and 561 nm (6.0%) lasers. Standard deconvolution was performed in Zen Black. For analysis of mitochondrial morphology in 3D and quantification of ATP synthase puncta from the obtained SIM images, individual cells were cropped in ImageJ (Fiji, NIH). Using Imaris 10.1.0, objects in separate channels were segmented and rendered in 3D with the Surfaces function. Segmentation setup included smoothing with Surfaces detail of 0.0986 µm, and background subtraction (Local Contrast) of 0.370 and 0.2 µm were used for TOMM20 and ATP Synthase channels, respectively. Machine Learning Segmentation was used for Hoechst channel with smoothing and Surfaces detail of 1 µm. Manual thresholding was used for TOMM20 and ATP Synthase channels. For ATP Synthase channel, Split touching Objects (Region Growing) was enabled with an Intensity Based Seed Points Diameter of 0.2 µm. A surfaces filter was then applied to select objects above 10 voxels. Object statistics from Imaris surfaces were then recorded in Microsoft Excel.

### Transmission electron microscopy (TEM)

BMDMs were plated at 1 x 10^6^ cells/mL (2 mL total volume) in a 35 mm dish with plastic cover slips and left to adhere overnight at 37°C in a de-humidified incubator (21% O_2_, 5% CO_2_). After stimulation, cells were washed with PBS and then fixed in fixation buffer (2% glutaraldehyde, 2% formaldehyde) in 50 mM sodium cacodylate pH 7.4 containing 2 mM calcium chloride for 2 h at room temperature. After 2 h, cells were kept in fixation buffer at 4°C overnight. Cells were subsequently washed three times with washing buffer (50 mM sodium cacodylate pH 7.4) and samples were osmicated (1% osmium tetroxide, 1.5% potassium ferricyanide and 50mM sodium cacodylate pH 7.4) for 3 days at 4°C. After washes with deionised water, samples were treated with 0.1% (w/v) thiocarbohydrazide in deionised water for 20 mins at room temperature in the dark. Samples were then washed in deionised water and osmicated a second time for 1 h at room temperature (2% osmium tetroxide in deionised water). After washes with deionised water, samples were blockstained with uranyl acetate (2% uranyl acetate in 50 mM maleate pH 5.5) for 3 days at 4°C. Then, samples were washed in deionised water and dehydrated in a graded series of ethanol (50%/70%/95%/100%/100% dry) and 100% dry acetonitrile, three times for each at least 5 mins. Samples were then infiltrated with a 50/50 mixture of 100% dry acetonitrile/Quetol resin (without BDMA) overnight, followed by three days in 100% Quetol (without BDMA). Then, samples were infiltrated for five days in 100% Quetol resin with BDMA exchanging the resin each day. The dishes were filled with resin to the rim, covered with a sheet of Aclar and cured at 60°C for three days. After curing, the Aclar sheets were removed, and small blocks were cut from the dishes using a hacksaw. Thin sections (∼70 nm) were prepared using an ultramicrotome (Leica Ultracut E). Resin blocks were oriented with the cell-side towards the knife and sections were collected on bare 300 mesh copper grids immediately when reaching the cell monolayer. Samples were then imaged with a Tecnai G2 TEM (FEI/ThermoFisher Scientific) run at 200 keV using a 20µm objective aperture to improve contrast. Images were acquired using an ORCA HR high resolution CCD camera (Advanced Microscopy Techniques Corp, Danvers USA). Mitochondrial morphology was assessed semi-quantitatively and mitochondrial length measured quantitatively using Fiji (NIH). At least 12 images per mouse per condition were analysed in each experiment.

### Proteomic analysis

BMDMs at 0.5 x 10^6^ cells/mL (5 mL total volume) were plated onto 6-cm dishes and left to adhere overnight at 37°C in a de-humidified incubator (21% O_2_, 5% CO_2_) before being treated as indicated. At the experimental endpoint, cells were washed with PBS on ice, scraped and centrifuged at 1500 rpm for 5 mins at 4°C and frozen at-80°C. Cell pellets were lysed, reduced and alkylated in 50 µl of 6 M Gu-HCl, 200 mM Tris-HCl pH 8.5, 10 mM TCEP, 15 mM chloroacetamide by probe sonication and heating to 95°C for 5 mins. Protein concentration was measured by a Bradford assay and initially digested with LysC (Wako) with an enzyme to substrate ratio of 1/200 for 4 h at 37 °C. Subsequently, the samples were diluted 10-fold with water and digested with porcine trypsin (Promega) at 37°C overnight. Samples were acidified to 1% TFA, cleared by centrifugation (16,000 g at RT) and approximately 20 µg of the sample was desalted using a Stage-tip. Eluted peptides were lyophilized, resuspended in 0.1% TFA/water and the peptide concentration was measured by A280 on a nanodrop instrument (Thermo). The sample was diluted to 2 µg/ 5 µl for subsequent analysis.

The tryptic peptides were analyzed on a Fusion Lumos mass spectrometer connected to an Ultimate Ultra3000 chromatography system (both Thermo Scientific, Germany) incorporating an autosampler. 2 µg of de-salted peptides were loaded onto a 50 cm emitter packed with 1.9 µm ReproSil-Pur 200 C18-AQ (Dr Maisch, Germany) using a RSLC-nano uHPLC systems connected to a Fusion Lumos mass spectrometer (both Thermo, UK). Peptides were separated by a 140 min linear gradient from 5% to 30% acetonitrile, 0.5% acetic acid. The mass spectrometer was operated in DIA mode, acquiring a MS 350-1650 Da at 120k resolution followed by MS/MS on 45 windows with 0.5 Da overlap (200-2000 Da) at 30k with a NCE setting of 27.

Raw files were analysed and quantified by searching against the Uniprot *Mus Musculus* data base using DIA-NN 1.8 (https://github.com/vdemichev/DiaNN). Library-free search was selected, and the precursor ion spectra were generated from the FASTA file using the deep learning option. Default settings were used throughout apart from using “Robust LC (high precision)”. In brief, Carbamidomethylation was specified as fixed modification while acetylation of protein N-termini was specified as variable. Peptide length was set to minimum 7 amino acids, precursor FDR was set to 1%. Subsequently, missing values were replaced by a normal distribution (1.8 π shifted with a distribution of 0.3 π) in order to allow the following statistical analysis. Protein-wise linear models combined with empirical Bayes statistics are used for the differential expression analyses. We use the Bioconductor package limma to carry out the analysis as previously described^72^. Heatmaps were generated using Morpheus software from the Broad Institute.

### RNA sequencing

BMDMs were plated at 0.5 x 10^6^ cells/mL (2 mL total volume) in a 6-well dish and left to adhere overnight at 37°C in a de-humidified incubator (21% O_2_, 5% CO_2_) before being treated as indicated. RNA isolation was carried using RNeasy^®^ Plus kit (74136, Qiagen) following manufacturer’s suggestions and eluted RNA was purified using RNA Clean & Concentrator Kits (Zymo Research). RNA-seq samples libraries were prepared by Cambridge Genomic Services (CGS) using TruSeq Stranded mRNA (Illumina) following the manufacturer’s description. For the sequencing, the NextSeq 75 cycle high output kit (Illumina) was used and samples spiked in with 1% PhiX. The samples were run using NextSeq 500 sequencer (Illumina). Differential Gene Expression Analysis was done using the counted reads and the R package edgeR version 3.26.5 (R version 3.6.1) for the pairwise comparisons.

### Metabolomic analysis

BMDMs were plated at 0.5 x 10^6^ cells/mL (1 mL total volume) in a 12-well dish and left to adhere overnight at 37°C in a de-humidified incubator (21% O_2_, 5% CO_2_), before being treated as indicated. For the stable isotope-assisted tracing experiments, glutamine free-DMEM (Gibco^TM^) was supplemented with U-^13^C-glutamine (Cambridge Isotope Laboratories) and replaced complete DMEM at the experimental start point. For the glucose-and glutamine-free experiments, DMEM without glucose and DMEM without glutamine (Gibco^TM^) were replaced at the experimental start point prior to being treated as indicated. Metabolite extraction buffer (MES) (methanol/acetonitrile/water, 50:30:20 v/v/v) was added (0.25 mL per 0.5 x 10^6^ cells) and samples were incubated for 15 mins on dry-ice. The resulting suspension was transferred to ice-cold microcentrifuge tubes. Samples were agitated for 20 mins at 4°C in a thermomixer and then incubated at - 20°C for 1 h. Samples were centrifuged at maximum speed for 10 min at 4°C. The supernatant was transferred into a new tube and centrifuged again at maximum speed for 10 min at 4°C. The supernatant was transferred to autosampler MS vials and stored at −80°C. Metabolite extracts were dried using an SC210A SpeedVac vacuum centrifuge (ThermoFisher Scientific) and reconstituted in an appropriate LC sample buffer, as detailed below. When necessary, further internal standards were added to the LC sample buffer: universal ^15^N^13^C amino acid mix, succinate ^13^C_4_; AMP ^15^N_2_^13^C_10_; ATP ^15^N_5_^13^C_10_; putrescine D8; dopamine D4. Internal standards were not included in the LC sample buffer for ^13^C labelling experiments. A Q Exactive Plus orbitrap coupled to a Vanquish Horizon ultra-high performance liquid chromatography system (ThermoFisher Scientific) was used for LC-MS analysis. Several customised LC methods were used to separate aqueous metabolites of interest as previously published^73^.

### HILIC method

Samples were reconstituted in 7:3 acetonitrile: water and analysed using a bridged ethylene hybrid (BEH) amide hydrophilic interaction liquid chromatography (HILIC) approach for the highly polar aqueous metabolites. The LC column used was the Acquity Premier BEH amide column (150 × 2.1 mm, 1.7 μm, Waters, cat 186009506). Mobile phase (A) was 100 mM ammonium carbonate and mobile phase (B) was acetonitrile, with 1:1 water:acetonitrile being used for the needle wash. The flow rate was 0.5 mL/min and the injection volume was 5 μL. The following linear gradient was used: 20% (A) for 1.5 mins, followed by an increase to 60% (A) over 3.5 mins, a hold at 60% (A) for 1 minute, a return to 20% (A) over 0.1 min and column re-equilibration for 3.9 mins. The total run time was 10 mins.

### C18pfp method

An ACE Excel C18-PFP column (150 x 2.1 mm, 2.0µm, Avantor, cat EXL-1010-1502U) was used to separate TCA cycle intermediates and amino acids. Samples were reconstituted in 10 mM ammonium acetate. For positive ion mode, mobile phase (A) was water with 10 mM ammonium formate and 0.1% formic acid and mobile phase (B) was acetonitrile with 0.1% formic acid. For negative ion mode, mobile phase (A) was water with 0.1% formic acid, and mobile phase (B) was acetonitrile with 0.1% formic acid. The flow rate was 0.5 mL/min and the injection volume was 3-3.5 μL. The needle wash used was 1:1 water:acetonitrile. The following gradient was used: 0% (B) for 1.6 mins, followed by a linear gradient to 30% (B) over 2.4 mins, a further increase to 90% (B) over 0.5 mins, a hold at 90% (B) for 0.5 mins, a return to 0% (B) over 0.1 min, and re-equilibration for 1.4 mins. The total run time was 6.5 mins.

### BEH C18 anion exchange (high mass) method

An Atlantis Premier NEH C18 AX column (Waters, cat 186009368) was used for the separation of higher mass species. Samples were reconstituted in 10 mM ammonium acetate prior to injection. Mobile phase (A) was 10 mM ammonium acetate and mobile phase (B) was 9:1 ACN:H20, 10 mM ammonium acetate, 0.1% ammonia. The flow rate was 0.4 mL/min and the injection volume was 5 μL. The needle wash used was 1:1 water:acetonitrile. The following gradient was used: 0% (B) for 1.6 minutes, followed by a linear gradient to 30% (B) over 1.9 minutes, a further increase to 90% (B) over 1 min, a hold at 90% B for 0.5 mins, a return to 0% B over 0.1 min, with re-equilibration for 2.4 mins. The total run time was 6.5 mins.

Source parameters used for the orbitrap were an auxiliary gas heater temperature of 438°C, a capillary temperature of 269°C, an ion spray voltage of 3.5 kV and a sheath gas, auxiliary gas and sweep gas of 53, 14 and 3 arbitrary units respectively with an S-lens radio frequency of 85%. A full scan of 500-1,000 m/z was used at a resolution of 70,000 ppm in positive ion mode.

To measure L-and D-enantiomers of 2-HG, diacetyl-L-tartaric anhydride-(DATAN, Merck, cat 336040050)-derivatised metabolite extracts from U-^13^C_-_glutamine labelling experiments were separated on an Acquity Premier HSS T3 column (1.8 µm, 2.1 x 100 mm, Waters, cat 186009468), as previously described^48^. The mobile phases were: (A) 1.5 mM ammonium formate (to pH 3.6 with formic acid) and (B) acetonitrile with 0.1% formic acid. The flow rate was 0.4 ml/min and the injection volume 5 µL. The column gradient used was: 3 to 5% (B) over 2.6 mins, followed by 5 to 95% (B) over 0.4 minutes, a hold at 95% (B) for 1.7 minutes, a return to 3% (B) over 1.3 mins and re-equilibration at 3% (B) for 6 mins, for a total run time of 12 mins.

Source parameters used for the Q Exactive orbitrap were identical to the C18pfp method. Parallel reaction monitoring (PRM) of the following 6 transitions was performed in negative ion mode using a collision energy of 20: 363.0569 (2-HG M+0 + DATAN) to 147.0299 (2-HG M+0); 364.0603 (2-HG M+1 + DATAN) to 148.0333 (2-HG M+1); 365.0636 (2-HG M+2 + DATAN) to 149.0366 (2-HG M+2); 366.0670 (2-HG M+3 + DATAN) to 150.0400 (2-HG M+3); 367.0703 (2-HG M+4 + DATAN) to 151.0433 (2-HG M+4); and 368.0737 (2-HG M+5 + DATAN) to 152.0467 (2-HG M+5).

LC-MS data were analysed with a targeted approach using ThermoFisher Scientific Xcalibur. Peak areas were normalised to an appropriate internal standard. Fractional incorporation (%) of individual isotopomers was calculated from the sum of all isotopomers. For L- and D- 2-HG, the m/z ratios of 147.0299 (2-HG M+0), 148.0333 (2-HG M+1), 149.0366 (2-HG M+2), 150.0400 (2-HG M+3), 151.0433 (2-HG M+4) and 152.0467 (2-HG M+5) were extracted in Xcalibur prior to integration of L- and D- peaks and subsequent calculation of fractional incorporation.

### Coenzyme Q (CoQ) extraction and analysis

BMDMs were plated at 0.5 x 10^6^ cells/mL (2 mL total volume) in a 6-well dish and left to adhere overnight at 37°C in a de-humidified incubator (21% O_2_, 5% CO_2_). The following day, cells were washed twice in cold PBS and then scraped into 200µL PBS, added to a mixture of 300 µL acidified methanol (0.1% (w/v) HCl) and 250µL hexane, and vortexed to extract CoQ. The hexane phase was separated by centrifugation at 17,000g for 5 mins. The hexane layer was transferred in mass spectrometry (MS) vials and dried under N_2_. The crude residue was resuspended in methanol containing ammonium formate (2 mM), overlaid with argon and analysed by LC-MS/MS using a I-class Acquity LC system attached to a Xevo TQ-S triple quadrupole mass spectrometer (Waters). Samples were stored in a refrigerated autosampler and 2 µL of sample was injected into a 15 µL flowthrough needle and separated by RP-HPLC at 45°C using an Acquity column (Waters). The isocratic mobile phase was ammonium formate (2 mM) in methanol used at 0.8mL/min over 5 mins. The mass spectrometer was operated in positive ion mode with multiple reaction monitoring. The following settings were used for electrospray ionisation in positive ion mode: capillary voltage – 1.7kV; cone voltage – 30V; ion source temperature – 100°C; collision energy – 22V. Nitrogen and argon were used as curtain and collision gases respectively. Transitions used for quantification were: UQ_9_, 812.9>197.2; UQ_9_H_2_, 814.9>197.2. Samples were quantified using MassLynx 4.1 software to determine the peak of area for UQ_9_ and UQ_9_H_2_.

### ATP, ADP and AMP measurements

BMDMs were plated at 0.5 x 10^6^ cells/mL (2 mL total volume) in a 6-well dish and left to adhere overnight at 37°C in a de-humidified incubator (21% O_2_, 5% CO_2_). The extraction buffer used was 4:1 MeOH: water with 5 μM ATP-^13^C (710695, Merck) as the internal standard (IS). Keeping the plate on ice, cells were quickly washed twice in PBS, 1 mL of extraction buffer was added and then scraped, transferred to a microcentrifuge tube and stored at-80°C for 1 h. A 0.2 μm PTFE filter was used to remove the protein in the sample and diluted 10-fold using the extraction buffer with IS prior to injection on a HPLC column (Shimadzu) mass spectrometer (Waters). An Atlantis Premier BEH Z-HILIC 1.7 μm, 2.1 x 150 mm (Waters) was used. Mobile phase A was 15 mM ammonium acetate with 0.1% ammonium and mobile phase B was 100% MeOH. The flow rate was 0.2 mL/min, and the injection volume was 5 µL. Peak areas were normalised to an appropriate internal standard. ATP, ADP and AMP concentrations in the samples were determined using a standard curve and the ratios assessed.

### Oxylipin analysis

BMDMs at 1 x 10^6^ cells/mL (2 mL total volume) were plated onto 6-cm dishes and left to adhere overnight at 37°C in a de-humidified incubator (21% O_2_, 5% CO_2_). Medium was replaced with phenol red-free DMEM (Gibco^TM^) before being treated as indicated. Cell culture supernatant was removed and stored at-80°C prior to extraction. Cell culture supernatant samples were spiked with 2.1-2.9ng 12-HETE-d8, 15-HETE-d8, LTB4- d4 and PGE_2_-d4 standards (Cayman Chemical) prior to extraction and downstream processing as previously described^74^. Lipids were extracted by adding a 2.5 mL solvent mixture (1M acetic acid/isopropanol/hexane; 2:30:30, v/v/v) to 1 mL of cells or cell culture supernatant in a glass extraction vial and vortexed for 1 min. 2.5 mL hexane was added to samples after vortexing for 1 min and tubes were centrifuged at 500g for 5 mins at 4°C to recover lipids in the upper hexane layer (aqueous phase), which was transferred to a new tube. Aqueous samples were re-extracted as above by addition of 2.5 mL hexane, and upper layers were combined. Lipid extraction from the lower aqueous layer was then completed according to the Bligh and Dyer technique using sequential additions of methanol, chloroform and water, and the lower layer was recovered following centrifugation as above and combined with the upper layers from the first stage of extraction. Solvent was dried under vacuum and lipid extract was reconstituted in 100 µL HPLC grade methanol. Lipids were separated by liquid chromatography using a gradient of 30-100% Acetonitrile: Methanol – 80:15+0.1% Acetic acid over 20 mins on an Eclipse Plus C18 Column (Agilent) and analysed on a Sciex QTRAP® 7500 LC-MS/MS system. Source conditions: TEM 475°C, IS-2500, GS1 40, GS2 60, CUR 40. Lipids were detected using MRM monitoring with the following parent to daughter ion transitions: PGD_2_ [M-H] – 351.2/271.1, PGE_2_ [M-H] – 351.2/271.1, 11-HETE [M-H] – 319.2/167.1, 8,9-DiHETrE [M-H] – 337.2/127.1, 14,15-DiHETrE [M-H] – 337.2/207.1, 17,18-DiHETE [M-H] – 335.2/247.1. Deuterated internal standards were monitored using parent to daughter ion transitions of 12-HETE-d8 [M-H] – 327.2/184.1, PGE_2_-d4 [M-H] – 355.2/275.1, 15- HETE-d8 [M-H] – 327.2/226.1, LTB4-d4 [M-H] – 339.2/197.1. Chromatographic peaks were integrated using Sciex OS 3.3.0 software (Sciex). Peaks were only selected when their intensity exceeded a 5:1 signal to noise ratio with at least 7 data points across the peak. The ratio of analyte peak areas to internal standard were taken and lipids quantified using a standard curve made up and run at the same time as the samples.

### Western Blot

BMDMs were plated at 0.5 x 10^6^ cells/mL (1 mL total volume) in a 12-well dish and left to adhere overnight at 37°C in a de-humidified incubator (21% O_2_, 5% CO_2_), before being treated as indicated. Cells were scraped in lysis buffer (Pierce™ RIPA Lysis and Extraction buffer, 89900, ThermoFisher Scientific) supplemented with 1X Protease Inhibitor Cocktail (ab271306, Abcam) and 5μL/mL Benzonase Nuclease (E1014-5KU, Merck). Total protein concentrations in samples were measured using a Pierce™ BCA Protein Assay Kit (23227, ThermoFisher Scientific) and diluted to the same concentration as required. Samples were denatured and reduced for SDS-PAGE in 4X Bolt™ LDS Sample Buffer (B0007, Invitrogen) and 10X Bolt™ Sample Reducing Agent (B0009, Invitrogen) at 95°C for 5 mins. Equal protein amounts were loaded and separated on Bolt™ 4-12% Bis-Tris Plus Mini Protein Gels 4-12% (NW04125BOX, Invitrogen). After SDS-PAGE, proteins were semi dry-transferred to nitrocellulose membranes (IB23001, ThermoFisher Scientific) for 12 mins at 20V. Membranes were blocked in 5% BSA or milk in TBS-Tween 0.1% for 1 hour at RT and subsequently incubated with primary antibodies diluted in 5% BSA or milk in TBS-Tween 0.1% overnight at 4°C. After washing membranes three times with TBS-Tween 0.1%, membranes were incubated with a secondary antibody solution in 5% BSA or milk for 30 mins at RT. Lastly, membranes were washed three times with TBS-Tween 0.1% and imaged using an Amersham Imager 680 and the SignalFire™ Plus ECL Reagents (6883, Cell Signalling). Membranes were exposed for different durations to ensure proper visualisation of all proteins. Blots were subsequently analysed using Image Lab software (Bio-Rad).

### Nitrite assay

BMDMs were plated at 0.5 x 10^6^ cells/mL (1 mL total volume) in a 12-well dish and left to adhere overnight at 37°C in a de-humidified incubator (21% O_2_, 5% CO_2_), before being treated as indicated. Nitric oxide production was assessed by measuring extracellular nitrite levels using the Griess assay (G2930, Promega) according to the manufacturer’s instructions. In brief, 50 μL extracellular media from each sample was placed in a 96-well plate in duplicate. An equal volume of sulfanilamide solution was added to each well and left for 5-10 mins at room temperature (RT), protected from light. Then 50 μL N-(1-naphthyl) ethylenediamine (NED) solution was added to each well and incubated for 5-10 mins at RT, protected from light. Absorbance was measured at 540 nm and nitrite levels were quantified against a standard curve made up using 0.1 M stock of sodium nitrite.

### Extracellular lactate measurements

BMDMs were plated at 0.5 x 10^6^ cells/mL (1 mL total volume) in a 12-well dish and left to adhere overnight at 37°C in a de-humidified incubator (21% O_2_, 5% CO_2_). Medium was changed prior to being treated as indicated. Extracellular lactate concentration in the cell culture medium was determined using a Lactate-Glo^TM^ Assay (J5021, Promega), according to the manufacturer’s instructions. In brief, 50 μL extracellular media from each sample was placed in a 96-well plate in duplicate. An equal volume of lactate detection reagent was added to each well and left to incubate for 60 mins at RT. Luminescence was subsequently recorded using a plate-reading luminometer.

### Digitonin fractionation

BMDMs were plated at 1 x 10^6^ cells/mL (2 mL total volume) in a 6-well dish and left to adhere overnight at 37°C in a de-humidified incubator (21% O_2_, 5% CO_2_) prior to indicated treatments. Following treatment, cells were subject to digitonin fractionation as previously described^30^. In brief, cells were washed once with room-temperature PBS, scraped on ice into ice-cold PBS and pelleted at 500 g for 5 mins at 4°C. The supernatant was discarded, and the pellet was resuspended in 400 μL of extraction buffer (150 mM NaCl, 50 mM HEPES pH 7.4, and 25 μg.ml^-1^ digitonin). Samples were rotated at 4°C for 10 mins before being centrifuged at 2,000g at 4°C for 5 mins. The resulting supernatant constituted the cytosolic fraction, from which RNA and DNA were isolated using the RNeasy® Plus kit (74136, Qiagen) or the DNeasy Blood & Tissue kit (69504, Qiagen). Alternatively, the cytosolic fraction was concentrated with Strataclean resin (Agilent) and analysed by western blot. The pellet containing membrane-bound organelles was lysed for analysis by western blotting. To detect mtRNA and mtDNA in the cytosol, qPCR was performed using primers specific for *mt-Nd1* and *mt-Co3* on cDNA reverse-transcribed from cytosolic RNA (mtRNA) and on DNA isolated from the cytosolic fraction (mtDNA).

### Olink target T48 mouse cytokine and chemokine panel

BMDMs at 0.5 x 10^6^ cells/mL (5 mL total volume) were plated onto 6-cm dishes and left to adhere overnight at 37°C in a de-humidified incubator (21% O_2_, 5% CO_2_). Medium was replaced with DMEM (Gibco^TM^) before being treated as indicated. Cell culture medium was removed, centrifuged at 1500 rpm for 5 mins to remove any cell debris before an aliquot was taken and transferred to a microcentrifuge tube and stored at-80°C. Proteins were measured using the Olink® Target 48 Cytokine Panel (Olink Proteomics AB, Uppsala, Sweden), which employs Proximity Extension Assay (PEA) technology to simultaneously analyse 43 analytes with only 1 μL of each sample. In this method, pairs of oligonucleotide-labelled antibody probes bind to their respective target proteins and when in close proximity, the oligonucleotides hybridise in a pair-wise manner. The addition of DNA polymerase triggers proximity-dependent DNA polymerisation, generating a unique PCR target sequence. The resulting DNA sequence is subsequently detected and quantified using a microfluidic real-time PCR instrument (Olink Q100 machine). Samples were run in the Stratified Medicine Core Laboratory (SMCL) NGS Hub, Department of Medical Genetics, University of Cambridge. Three internal controls are added to each sample, the incubation control, the extension control and the detection control. The extension control is used for the data normalisation of each sample but is not used as a quality control measure. The incubation control and the detection control are used to monitor the quality of assay performance, as well as the performance of individual samples. Three external controls are included in the kit and analysed on each sample plate, negative control, sample control and calibrator. Negative controls are analysed in duplicates and sample control and calibrator are analysed in triplicates on each plate. The calibrator is used for data normalisation and both calibrator and sample control are used to monitor the quality of assay performance. Only data from runs that meet these quality control criteria are reported. Data is provided as normalised protein expression (NPX). For more detailed information, see panel specific validation data and the Olink NPX Signature manual available at the Olink website (www.olink.com).

### Enzyme-linked immunosorbent assay (ELISA)

BMDMs were plated at 0.5 x 10^6^ cells/mL (1 mL total volume) in a 12-well dish and left to adhere overnight at 37°C in a de-humidified incubator (21% O_2_, 5% CO_2_) prior to being treated as indicated. Concentrations of secreted IFN-β, IL-6, IL-1β and TNF-α were determined using the corresponding ELISA kits (IFN-β DY8234, IL-6 DY406, IL-1β DY401, and TNF-α DY410). In brief, MaxiSorp Nunc-Immuno 96-well plates (439454, ThermoFisher) were incubated overnight at RT with a capture antibody. After three washes with 0.05% Tween® 20 (P1379, Sigma) in PBS, the plates were blocked for an hour with 1% BSA in PBS. Incubation with sample or standards was performed at 4°C overnight after three washes with 1% BSA in PBS. Experiments were conducted using two technical replicates of 50 μL of culture media. For IL-6 and TNF-α, the culture media was diluted in reagent diluent 1:2 and 1:5, respectively. Subsequently, plates were washed and incubated with the appropriate detection antibody for two hours at RT. After washing, a working solution of 50 μL streptavidin-HRP was added to the plates for 20 mins and the plate was covered from direct light. After washing, 50 μL substrate solution (1:1 mixture of H_2_O_2_ and tetramethylbenzidine (R&D Systems) was added to all wells and left to incubate for approximately 20 mins. The reaction was stopped using 25 μL 2 M H_2_SO^4^, and optical density measurements were collected at 450 nm on a SpectraMax Plus 384 microplate reader. Concentrations were calculated using the corresponding standard curve after accounting for the dilution of the sample in the assay when necessary.

### RT-qPCR

BMDMs were plated at 0.5 x 10^6^ cells/mL (1 mL total volume) in a 12-well dish and left to adhere overnight at 37°C in a de-humidified incubator (21% O_2_, 5% CO_2_) prior to being treated as indicated. mRNA expression was quantified by quantitative RT-qPCR. RNA extraction was performed as described in the manufacturer’s protocol (RNeasy^TM^ Plus Mini Kits 74136, Qiagen). cDNA was synthesized with the High-Capacity cDNA Reverse Transcription kit (ThermoFisher Scientific). The reaction was performed in a MicroAmp optical 96-well reaction plate (N8010560, Applied Biosystems) in a 96-well QuantStudio^TM^ 3 PCR machine with PowerUp™ SYBR™ Green Master Mix (A25741, Applied Biosystems) using primers listed in Table S1. *Rps18* was used as normalisation control.

### *Salmonella typhimurium* (STM) infection assay

Cells were primed for 4 h with LPS from *E. coli* serotype EH100 (Alexis) at a concentration of 100 ng/mL. *Salmonella enterica* serovar Typhimurium (*S.* Typhimurium) strain SL1344 was grown in low-salt lysogeny broth (LB) medium (ThermoFisher Scientific) at 37°C in an orbital shaker. Overnight *Salmonella* cultures were sub-cultured 1:10 in fresh LB and grown until mid-exponential phase (OD_600_ = 0.8–1.2). Prior to infection, bacteria were pelleted by centrifugation (9000 rpm, 1.5 mins), washed, and resuspended in Opti-MEM (Gibco). Bacteria were added to confluent cells in 96-well plates (BMDMs, 5 × 10^4^ cells/well) at different multiplicities of infection (MOIs), as described in the figure legends. Plates were then incubated for 30 mins at 37°C. Noninternalized bacteria were then removed by washing cells with prewarmed medium. Cells were incubated with Opti-MEM containing 10 μg/ml of gentamycin to eliminate any remaining extracellular bacteria. At specified time points postinfection (p.i.), cells were either processed for Colony-Forming Unit (CFU) analysis, Lactate Dehydrogenase Cytotoxicity assay (LDH), or ELISA. To determine CFU, infected cells were gently washed with PBS and lysed with water containing 0.2% Triton X-100 at the indicated time points. Bacteria were then serially diluted and plated onto LB agar. Cell death was quantified by measuring LDH release into the supernatant, using the LDH cytotoxicity detection kit (CytoTox 96® Non-Radioactive Cytotoxicity Assay, Promega). To normalize for spontaneous cell lysis, the percentage of cell death was calculated as follows: (LDH_sample_ – LDH_negative control_)/(LDH_positive control_ – LDH_negative control_) × 100. The level of IL-1β release was measured by ELISA as previously described.

### Endotoxemia model

Both male and female WT and m.5019A>G mice (8-12 weeks old) were used for these experiments. Mice were injected intraperitoneally with PBS or Ultrapure LPS from *Escherichia coli* O55:B5 (2.5 mg/kg, Sigma). After 2 h, mice were euthanised in a CO_2_ chamber and blood was collected from the Vena Cava. Blood was centrifuged at 1,500 rpm for 10 mins at 4°C and the serum was collected for analysis using an Olink target T48 mouse cytokine and chemokine panel and an IFN-β ELISA, as previously described. The pain response was also assessed using the NC3Rs grimace scale for mice, with 0 = not present, 1 = moderately present and 2 = obviously present. A behavioural analysis of the mice post-injection with PBS or LPS was performed to generate a sepsis score, using a method adapted from^75^. In brief, the movement behaviour, flight reaction and body position were assessed and assigned a score in a blinded manner from video recordings. This also included an in-person assessment of the pain response as a parameter using the NC3Rs grimace scale for mice. The sum of these scores constituted the sepsis score.

### Quantification and statistical analysis

Details of all statistical analyses performed can be found in the figure legends. Data were expressed as mean ± standard error of the mean (s.e.m) or standard deviation (s.d.), unless stated otherwise. Graphpad Prism 10.2.3 was used to calculate statistics in plots using appropriate statistical tests depending on the data including two-tailed unpaired t test, one-way ANOVA and multiple t tests. Adjusted p values were assessed using appropriate correction methods, such as Tukey and Holm-Sidak tests. Scaling of features (metabolites or proteins) was used for heatmap generation from metabolomics. For proteomics data, protein signal intensity was converted to a log_2_ scale and biological replicates were grouped by experimental condition. Protein-wise linear models combined with empirical Bayes statistics were used for the differential expression analyses. The Bioconductor package limma was used to carry out the analysis using an R based online tool^72^. Data were visualised using a Volcano plot, which shows the log_2_ fold change (FC) on the x axis and the-log_10_ adjusted p value on the y axis. The cut-off for significant hits in the proteomics, Olink mouse T48 panel and RNA seq were set to FDR < 0.05. Overrepresentation analysis (ORA) of differentially abundant proteins and differentially expressed genes were determined using ShinyGO^76^. GSEA analysis of RNA seq was performed using the Broad Institutes GSEA 4.1.0^77^. Sample sizes were determined on the basis of previous experiments using similar methodologies. All depicted data points are biological replicates taken from distinct samples (mice), unless stated otherwise. Each figure consists of a minimum of 3 biological replicates, as indicated in the figure legends, from multiple independent experiments, unless stated otherwise.

## Acknowledgements

We thank Alexandra Karcanias, Julien Bauer and Stephanie Wenlock of Cambridge Genomic Services (CGS), Department of Pathology, University of Cambridge for RNA sequencing and bioinformatic analysis services. We thank the Stratified Medicine Core Laboratory (SMCL) NGS Hub, Department of Medical Genetics, University of Cambridge, for the Olink Services. We thank the Cambridge Advanced Imaging Centre (CAIC), University of Cambridge for the assistance with Transmission Electron Microscopy (TEM). We thank Reiner Schulte and the staff of the Cambridge Institute for Medical Research (CIMR) flow cytometry core facility for training and support for flow cytometry analysis. We thank Alice Sowton of the Kunji lab for Oroboros training and Roy Chowdhury for microscopy training in the Medical Research Council (MRC) Mitochondrial Biology Unit (MRC MBU). We thank the Murphy lab of the MRC MBU for helpful discussions. This project was supported by funding to Dr. Dylan G. Ryan from the MRC (MC_UU_00028) and a Wellcome Trust-Academy Medical of Sciences (AMS) springboard grant (G123514).

## Author contributions

**D.G.R** conceptualised the project. **E.M** and **D.G.R** were the lead experimentalists, designed all experiments, analysed and visualised the data and co-wrote the paper with input from all authors. **S.P.B** and **P.F.C** assisted with pyrosequencing, mtDNA copy number assessment and provided important intellectual input and assistance with the m.5019A>G mouse model. **A.M.C, C.S.Y** and **M.P.M** assisted with targeted LC-MS analysis for CoQ and ATP, ADP and AMP. **R.J.S** and **A.K** assisted with metabolomics and stable isotope-assisted tracing. **K.T, Y.M.K** and **E.M** assisted with mouse breeding, colony management, tissue acquisition and *in vivo* experiments using the m.5019A>G mice. **A.M.C** assisted with blinded sepsis scoring. **D.M.W** assisted with immunofluorescence, super-resolution microscopy and image analysis. **M.D** and **C.E.B** assisted with the *Salmonella typhimurium* infection experiments. **V.J.T** and **V.B.O.D** assisted with the oxylipin profiling. **R.K.** assisted with metabolomic sample preparation and Griess assay. **C.A.P** and **M.M** assisted with ^35^S-methionine labelling experiments. **J.B.S** developed the m.5019A>G mouse model and provided input.

**A.v.K** assisted with the proteomics. **M.P.M, M.M, V.B.O.D, C.E.B, A.K**, **P.C and A.v.K** oversaw a portion of the research programme. **D.G.R** obtained the funding, lead the study and oversaw the research programme.

## Figure legends

**Extended data figure 1.**
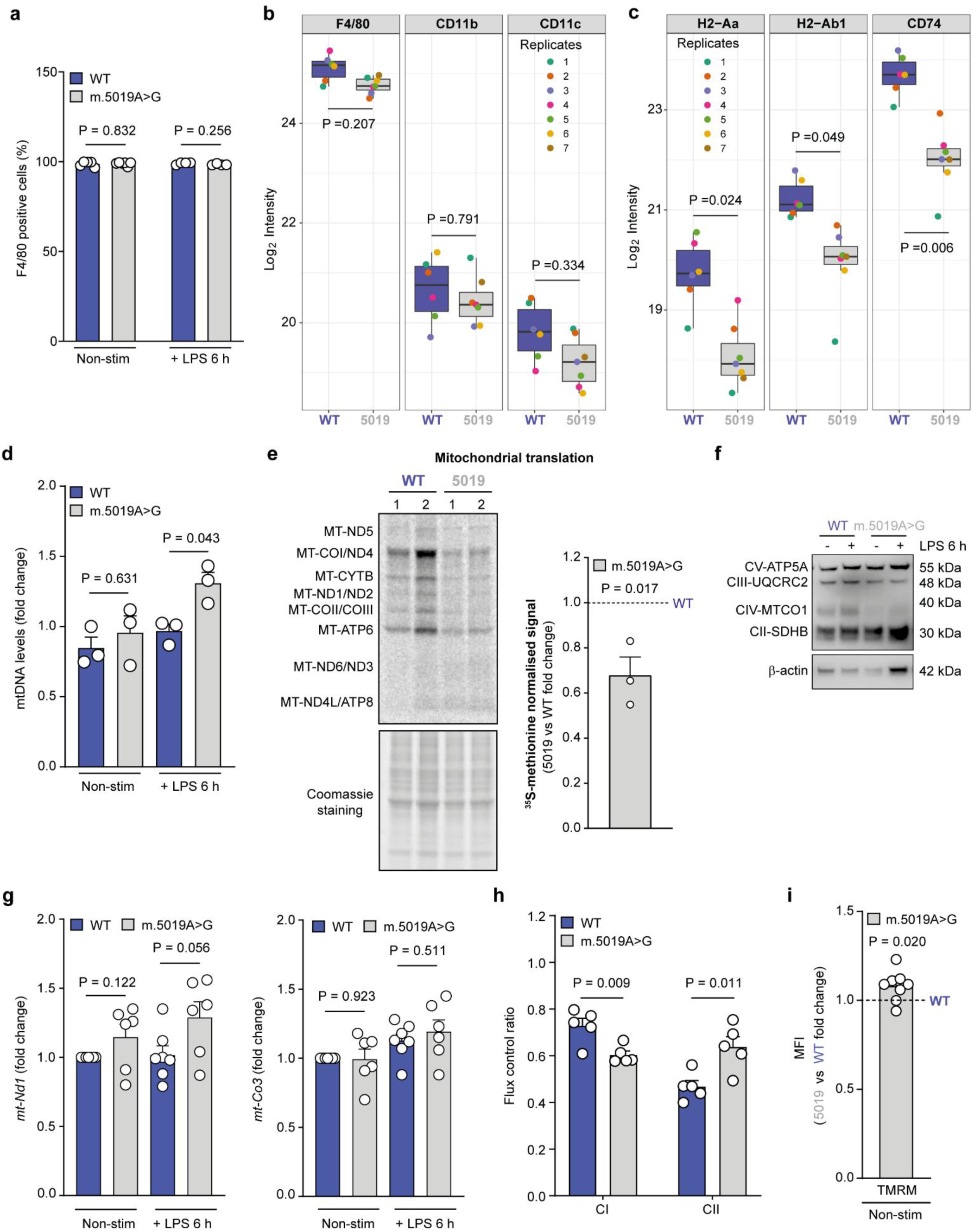
Reduced mitochondrial respiration in primary macrophages with a heteroplasmic m.5019A>G mtDNA mutation **a,** F4/80 cell surface staining of non-stim and LPS-stimulated WT and m.5019A>G BMDMs (*n* = 5; LPS 6 h). Proteomic analysis of (**b**) F4/80, CD11b and CD11c and (**c**) H2-Aa, H2-Ab1 and CD74 in non-stim WT and m.5019A>G (*n* = 6-7) BMDMs. **d**, mtDNA levels in non-stim and LPS-stimulated WT and m.5019A>G BMDMs (*n* = 3; LPS 6 h). **e**, ^35^S-methionine labelling and quantification of mitochondrial proteins in non-stim WT and m.5019A>G BMDMs (*n* = 3). **f**, CV-ATP5A, CIII-UQCRC2, CIV-MT-COI and CII-SDHB protein levels in non-stim and LPS-stimulated WT and m.5019A>G BMDMs (*n* = 2; LPS 6 h). Representative blot shown. **g**, *mt-Nd1* and *mt-Co3* expression in non-stim and LPS-stimulated WT and m.5019A>G BMDMs (*n* = 6; LPS 6 h). **h**, Complex I (CI) and CII flux control ratio in non-stim WT and m.5019A>G (*n* = 5). **i**, TMRM staining in non-stim m.5019A>G vs WT BMDMs (*n* = 8). Data are mean ± s.e.m. *P* values calculated using two-tailed Student’s t-test for two group comparisons.

**Extended data figure 2.**
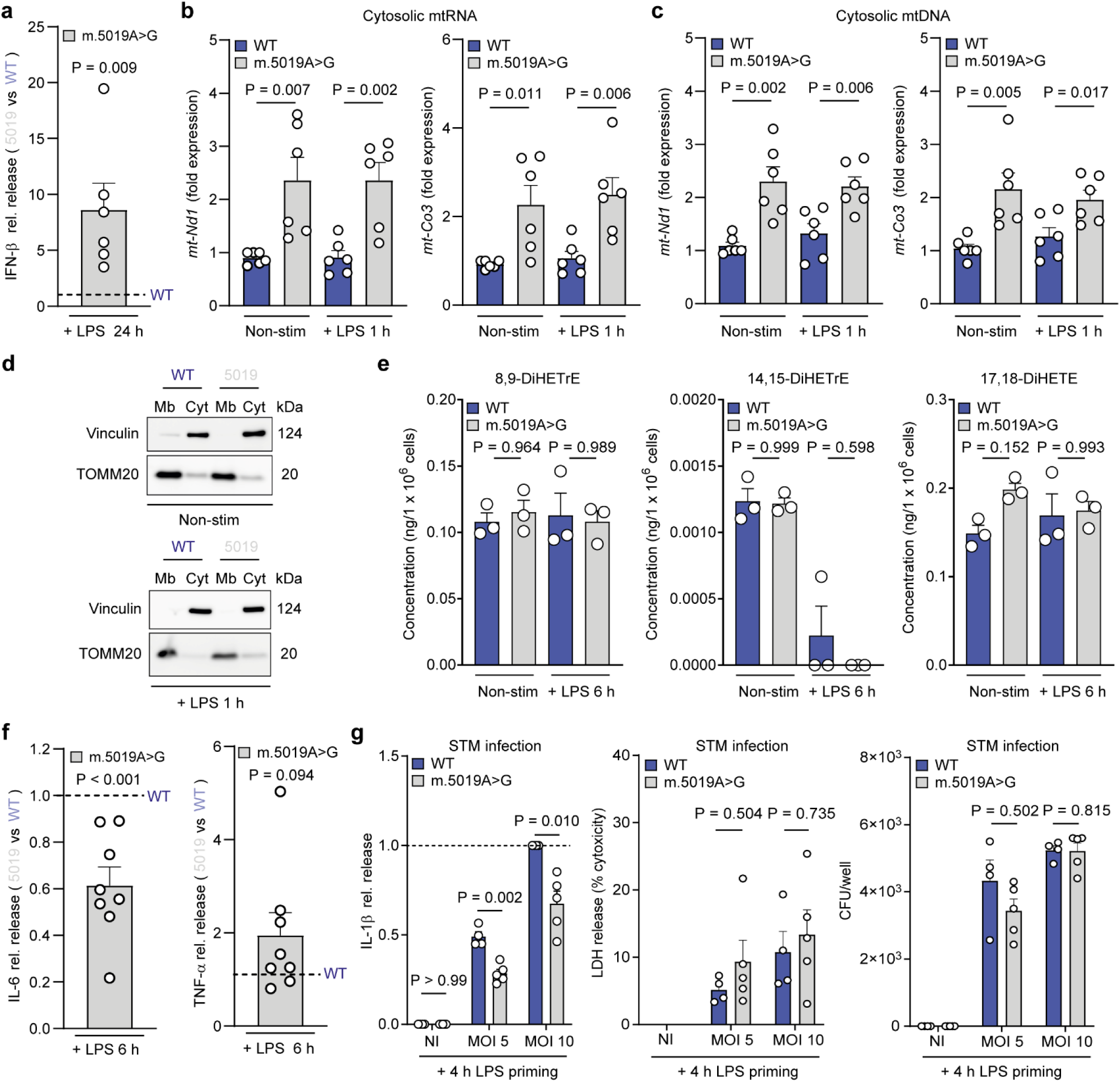
Inflammatory cytokine and oxylipin production are disrupted in m.5019A>G primary macrophages **a,** IFN-β release in LPS-stimulated m.5019A>G vs WT BMDMs (*n* = 6; LPS 24 h). *mt-Nd1* and *mt-Co3* (**b**) RNA and (**c**) DNA in cytosolic fraction of non-stim and LPS-stimulated WT and m.5019A>G BMDMs (*n* = 6; LPS 1 h). **d**, Vinculin and TOMM20 levels in membrane (Mb) and cytosolic (Cyt) fraction of non-stim and LPS-stimulated WT and m.5019A>G BMDMs (LPS 1 h; representative blot shown). **e,** Oxylipin profiling of CCM in non-stim and LPS-stimulated m.5019A>G vs WT BMDMs (*n* = 3; LPS 6 h). **f**, IL-6 and (**c**) TNF-α release in LPS-stimulated m.5019A>G vs WT BMDMs (*n* = 8; LPS 6 h). **g**, IL-1β release (30 mins), LDH release (30 mins) and bacterial burden (colony forming units (CFU); 1.5 h) following STM infection (multiplicity of infection (MOI) 5 and 10) of LPS-primed (4 h) WT and m.5019A>G BMDMs (*n* = 4-5). Data are mean ± s.e.m. *P* values calculated using two-tailed Student’s t-test for two group comparisons, multiple unpaired t-tests corrected for multiple comparisons using Holm-Sidak method.

**Extended data figure 3.**
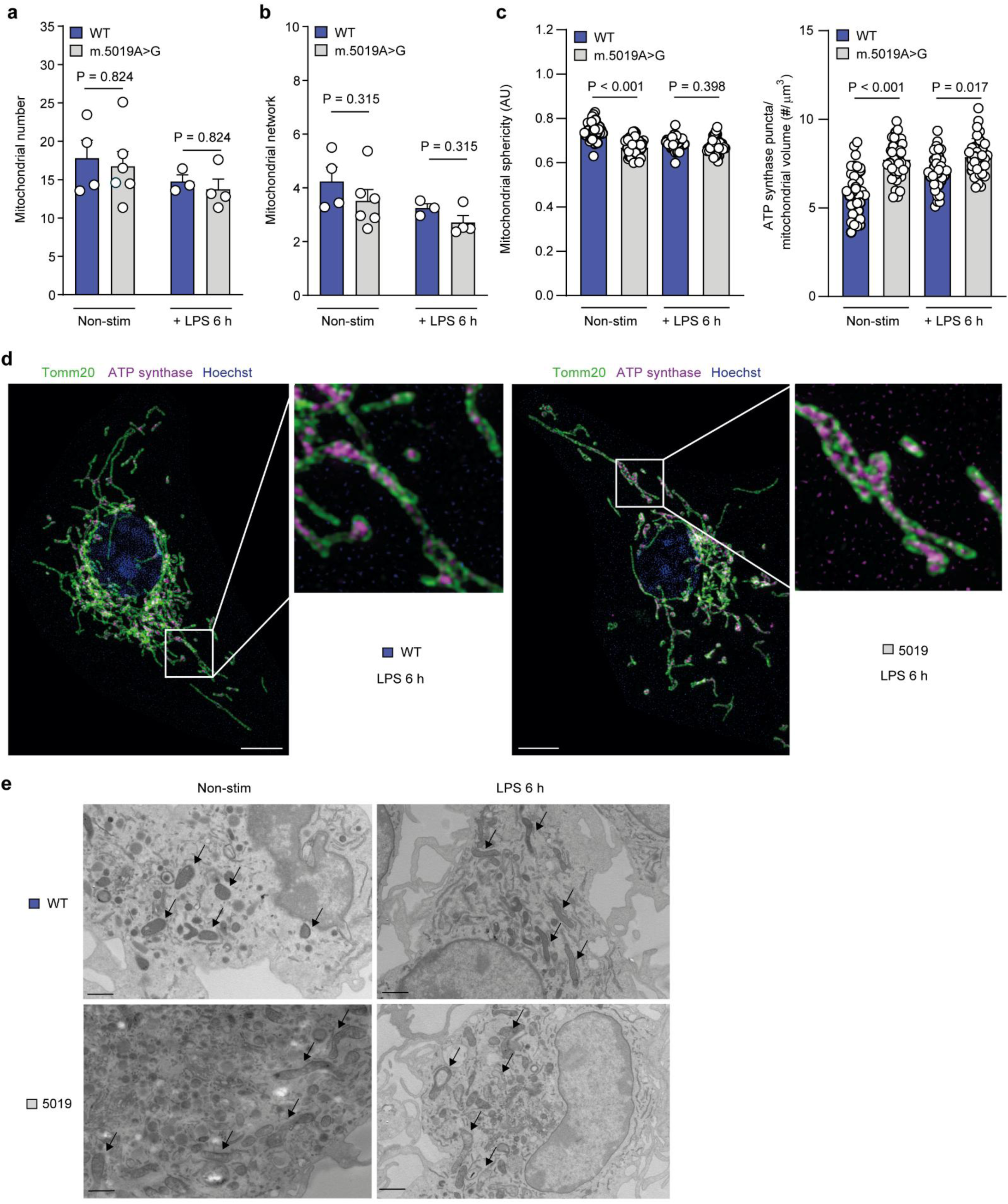
Increased mitochondrial elongation and disrupted cristae architecture in m.5019A>G primary macrophages **a-b**, Morphology analysis of immunofluorescence staining for Cyt c and TOMM20 coupled to confocal microscopy in non-stim (*n* = 4-6) and LPS-stimulated (*n* = 3-4; LPS 6 h) WT and m.5019A>G BMDMs. **c**, Morphology and ATP synthase puncta analysis of super-resolution microscopy images in non-stim and LPS-stimulated WT and m.5019A>G BMDMs (*n* = 2-3; LPS 6 h). **d**, Immunofluorescence staining of TOMM20 and ATP synthase coupled to super-resolution microscopy in LPS-stimulated WT and m.5019A>G BMDMs. Scale bar 5 μm. **e**, Transmission electron microscopy (TEM) of non-stim and LPS-stimulated WT and m.5019A>G BMDMs (*n* = 3; LPS 6 h). Scale bar 0.5 μm. Arrows indicate mitochondria. Representative images are shown. Data are mean ± s.e.m or ± s.d. *P* values calculated using two-tailed Student’s t-test for two group comparisons or one-way ANOVA corrected for multiple comparisons using the Kruskal-Wallis method.

**Extended data figure 4.**
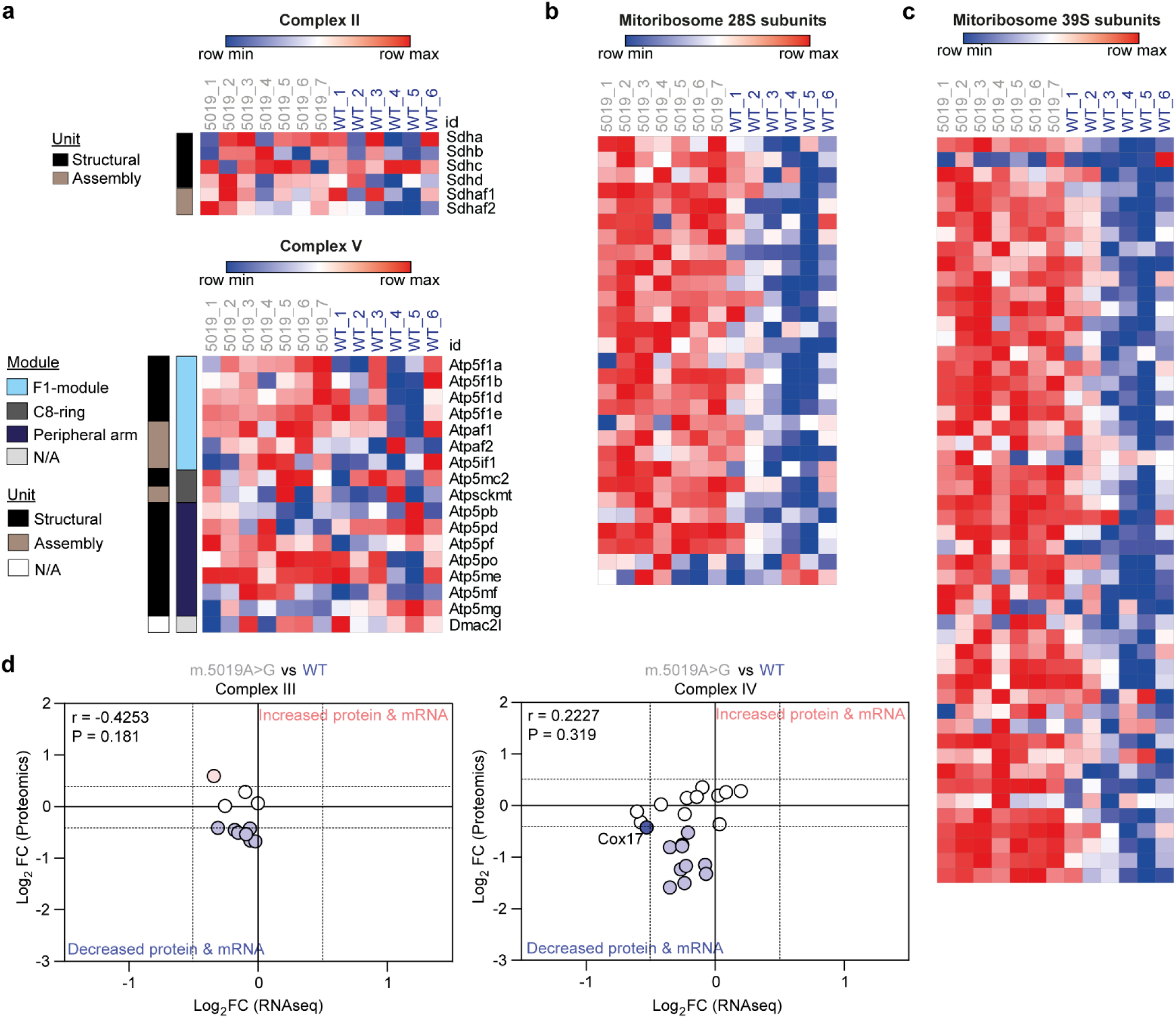
Decreased abundance of nuclear-encoded structural subunits of Complex I, III and IV in m.5019A>G primary macrophages Heatmap of (**a**) CII and CV (**b**) mitoribosome 28S and (**c**) mitoribosome 39S subunits in non-stim WT and m.5019A>G BMDMs (*n* = 6-7). **d**, Comparison of Log_2_FC values of CIII and CIV subunits from proteomics and RNA sequencing data with Pearson correlation and statistical analysis applied.

**Extended data figure 5.**
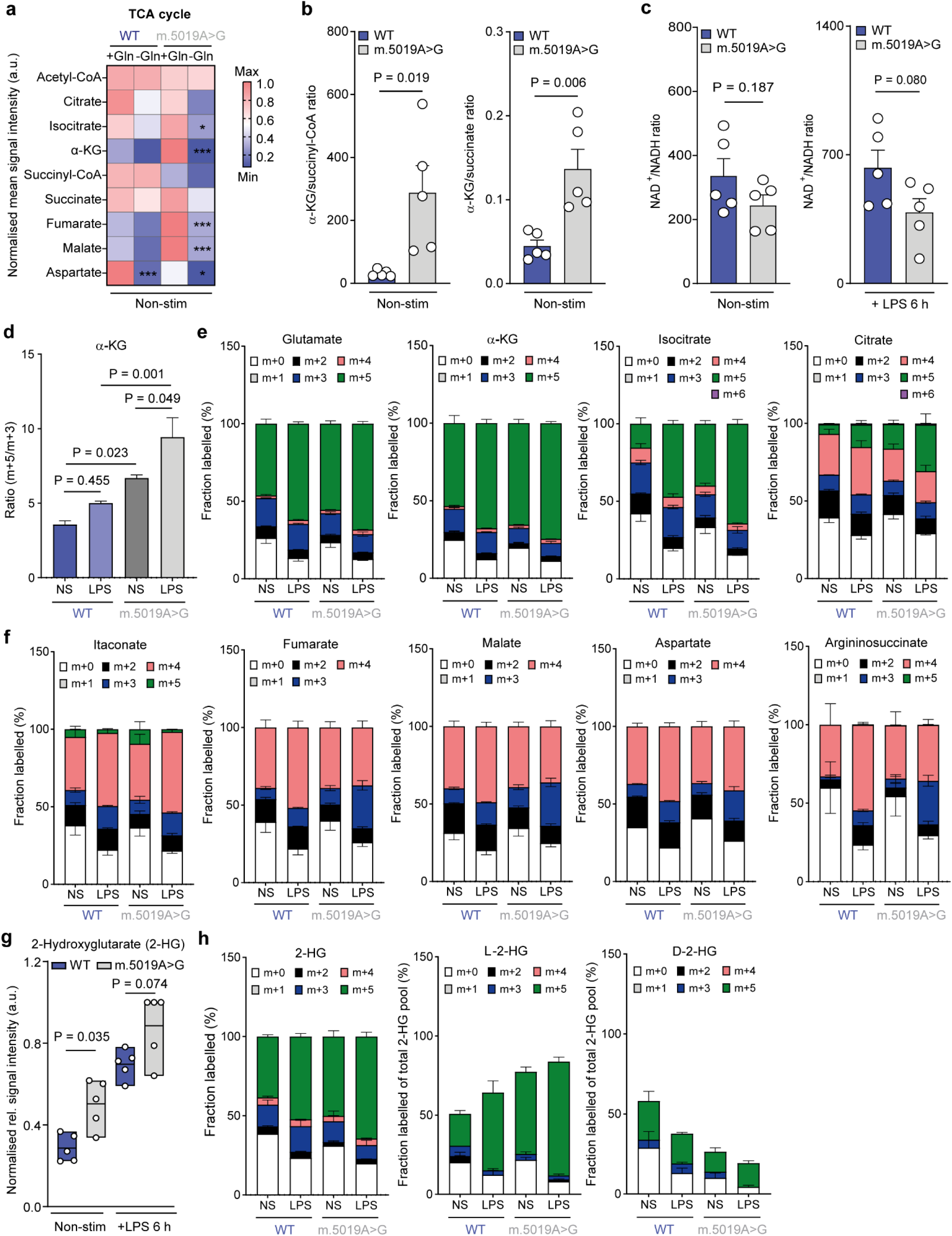
TCA cycle remodelling and increased nitric oxide production in m.5019A>G primary macrophages **a**, Heatmap comparing metabolite levels in non-stim WT and m.5019A>G BMDMs in the presence or absence of glutamine (Gln) (*n* = 3). **b**, α-KG/succinyl-CoA and α-KG/succinate ratio in non-stim WT and m.5019A>G BMDMs (*n* = 5). **c**, NAD^+^/NADH ratio in non-stim and LPS-stimulated WT and m.5019A>G BMDMs (*n* = 5; LPS 6 h). **d**, m+5/m+3 ratio of α-KG in non-stim and LPS-stimulated WT and m.5019A>G BMDMs (*n* = 5; LPS 6 h). **e**-**f**, Total isotopologue distribution (% fraction labelling) of TCA cycle metabolites, itaconate and aspartate from U-^13^C-glutamine tracing in non-stim and LPS-stimulated WT and m.5019A>G BMDMs (*n* = 5; LPS 6 h). **g**, 2-hydroxyglutarate (2-HG) levels in in non-stim and LPS-stimulated WT and m.5019A>G BMDMs (*n* = 5; LPS 6 h). **h**, Total isotopologue distribution (% fraction labelling) of 2-HG (*n* = 5), L-2-HG (*n* = 3) and D-2-HG (*n* = 3) from U-^13^C-glutamine tracing in non-stim and LPS-stimulated WT and m.5019A>G BMDMs (LPS 6 h). Data are mean ± s.e.m. *P* values calculated using two-tailed Student’s t-test for two group comparisons or multiple unpaired t-tests corrected for multiple comparisons using Holm-Sidak method or one-way ANOVA corrected for multiple comparisons using Tukey method. **\*\*\*** *P* < 0.001 **\*\*** *P* < 0.01 **\*** *P* < 0.05.

**Extended data figure 6.**
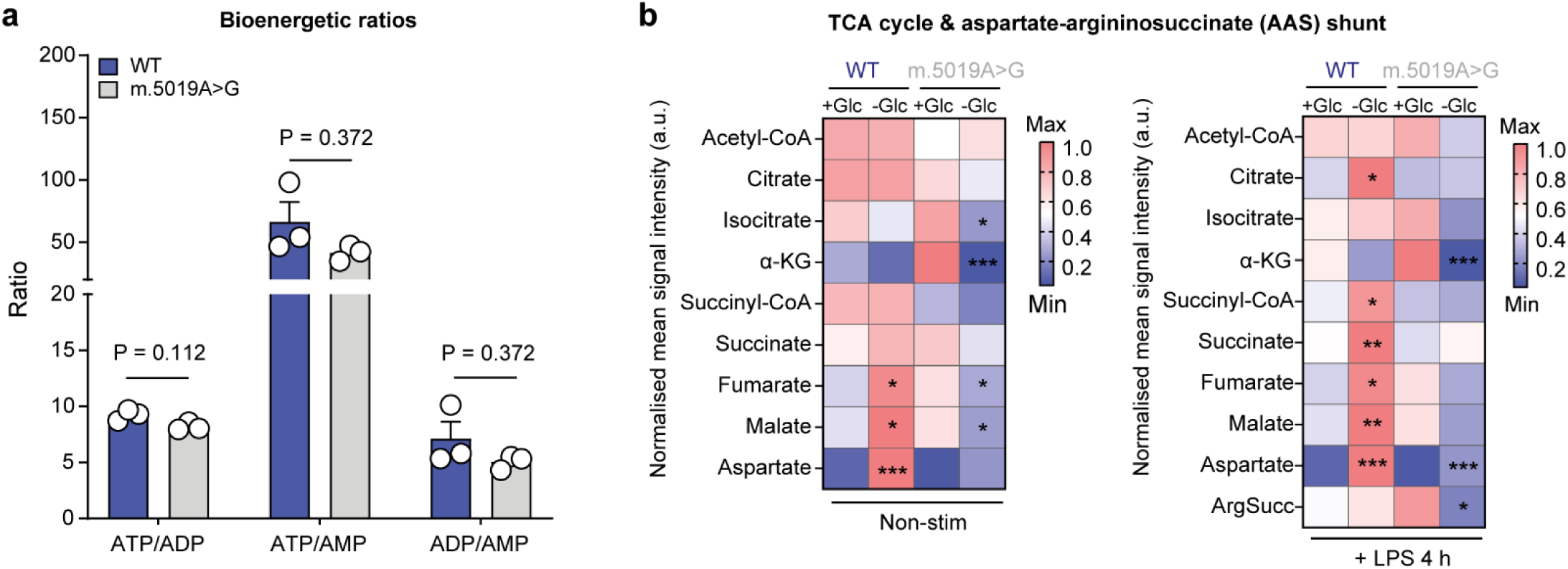
Increased aerobic glycolysis and glutathione in m.5019A>G primary macrophages **a,** ATP/ADP, ATP/AMP and ADP/AMP ratios in non-stim WT and m.5019A>G BMDMs (*n* = 3). **b**, Heatmap comparing TCA cycle and AAS metabolite levels in non-stim and LPS-stimulated WT and m.5019A>G BMDMs in the presence or absence of glucose (Glc) (*n* = 3; LPS 4 h). Data are mean ± s.e.m. *P* values calculated using two-tailed Student’s t-test for two group comparisons or multiple unpaired t-tests corrected for multiple comparisons using Holm-Sidak method. **\*\*\*** *P* < 0.001 **\*\*** *P* < 0.01 **\*** *P* < 0.05.

---

**Table S1.**
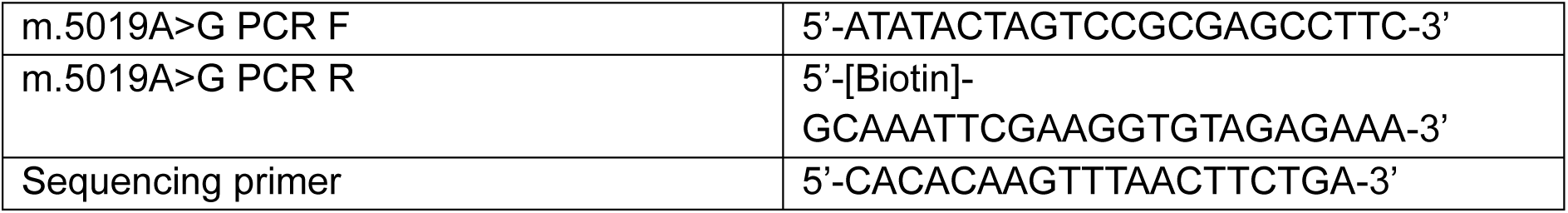
Pyrosequencing primers.

**Table S2.**
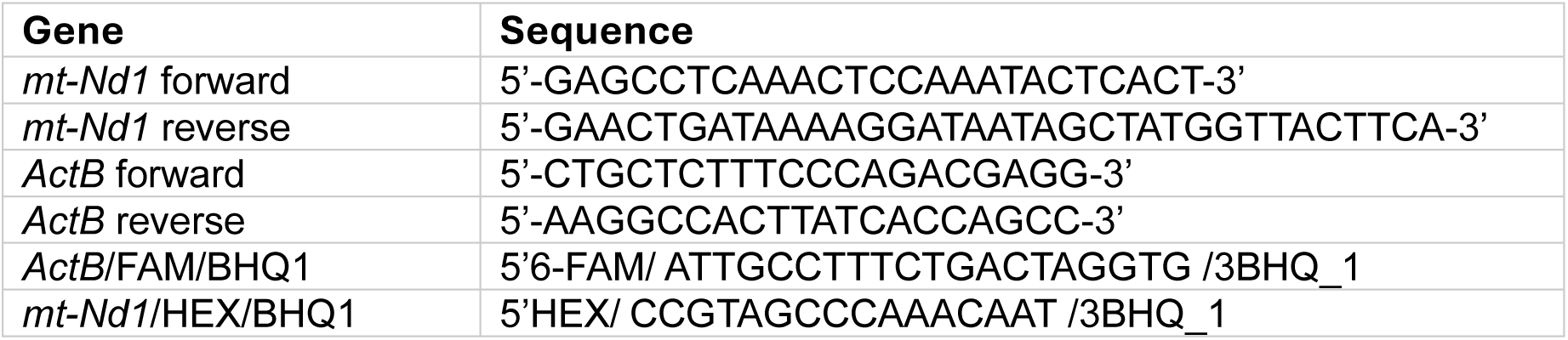
ddPCR primers.

**Table S3.**
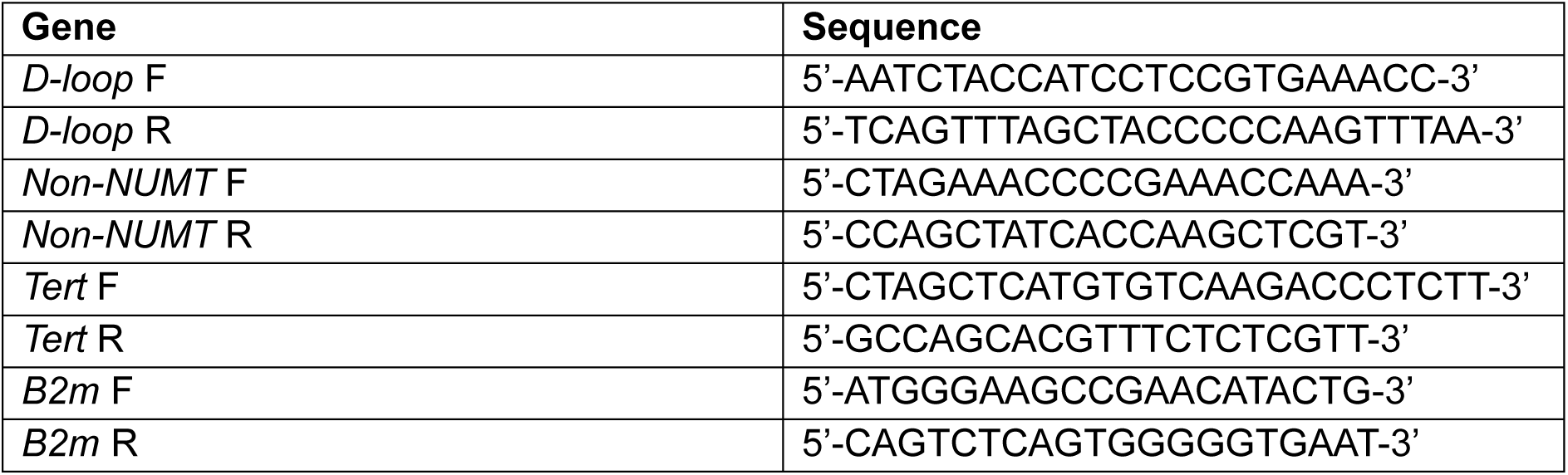
qPCR primers.

**Table S4.** Antibodies.

| Antibody | Source | Identifier |
| --- | --- | --- |
| Pro-IL-1β | Cell Signalling Technology | 12507; RRID: AB_2721117 |
| COX2 | Cell Signalling Technology | 12282S; RRID: AB_2571729 |
| iNOS | Cell Signalling Technology | 13120S; RRID: AB_2687529 |
| Total OXPHOS | Abcam | 110413; RRID: AB_2629281 |
| β-actin | Cell Signalling Technology | 4970; RRID: AB_2223172 |
| Cytochrome c | BD Biosciences | 556432; RRID: AB_396416 |
| TOMM20 | Proteintech | 11802-1-AP; RRID: AB_2207530 |
| Vinculin | Cell Signalling Technology | 13901; RRID: AB_2728768 |
| Anti-rabbit IgG, HRP-linked | Cell Signalling Technology | 7074 |

**Table S4.**
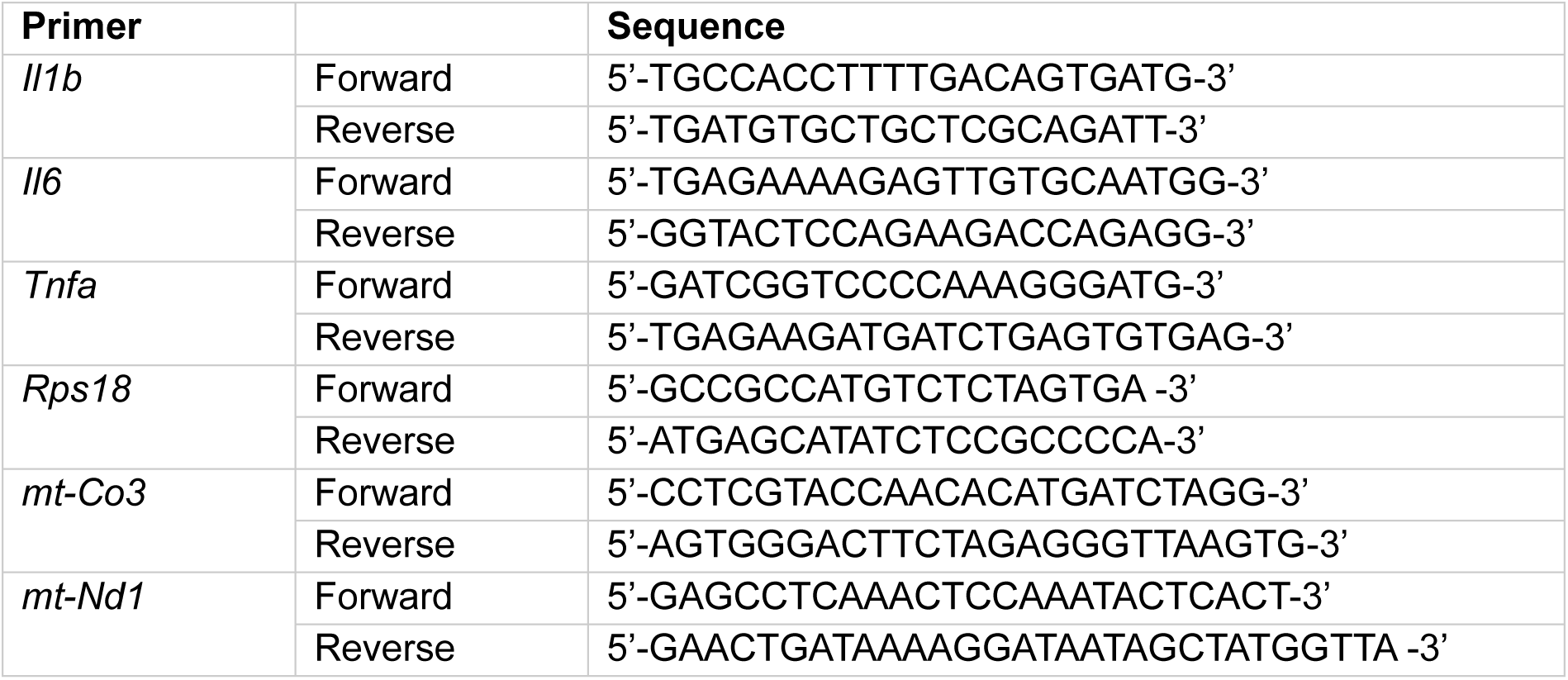
RT-qPCR primers.

## References

1. Y, L., et al. Elevated type I interferon responses potentiate metabolic dysfunction, inflammation, and accelerated aging in mtDNA mutator mice. Sci. Adv. 7, (2021).

2. Stokes, J. C., et al. Leukocytes mediate disease pathogenesis in the *Ndufs4*(KO) mouse model of Leigh syndrome. JCI Insight 7, (2022).

3. VanPortfliet, J. J. et al. Caspase-11 drives macrophage hyperinflammation in models of Polg-related mitochondrial disease. bioRxiv 2024.05.11.593693 (2024) doi:10.1101/2024.05.11.593693.

4. Jin, Z., Wei, W., Yang, M., Du, Y. & Wan, Y. Mitochondrial Complex I Activity Suppresses Inflammation and Enhances Bone Resorption by Tipping the Balance of Macrophage-Osteoclast Polarization. Cell Metab. 20, 483–498 (2014).

5. Chen, H. et al. Mitochondrial fusion is required for mtDNA stability in skeletal muscle and tolerance of mtDNA mutations. Cell 141, 280–289 (2010).

6. Hanaford, A. R. et al. Peripheral macrophages drive CNS disease in the Ndufs4(−/−) model of Leigh syndrome. Brain Pathol. 33, e13192 (2023).

7. Burr, S. P. et al. Cell lineage-specific mitochondrial resilience during mammalian organogenesis. Cell 186, 1212–1229.e21 (2023).

8. Hanaford, A. & Johnson, S. C. The immune system as a driver of mitochondrial disease pathogenesis: a review of evidence. Orphanet J. Rare Dis. 17, 335 (2022).

9. Marques, E., Kramer, R. & Ryan, D. G. Multifaceted mitochondria in innate immunity. Npj Metab. Health Dis. 2, 1–15 (2024).

10. Ryan, D. G., Frezza, C. & O’Neill, L. A. J. TCA cycle signalling and the evolution of eukaryotes. Curr. Opin. Biotechnol. 68, 72–88 (2020).

11. Inigo, M., Deja, S. & Burgess, S. C. Ins and Outs of the TCA Cycle: The Central Role of Anaplerosis. Annu. Rev. Nutr. 41, 19–47 (2021).

12. Stewart, J. B. & Chinnery, P. F. The dynamics of mitochondrial DNA heteroplasmy: implications for human health and disease. Nat. Rev. Genet. 16, 530–542 (2015).

13. Gorman, G. S. et al. Prevalence of nuclear and mitochondrial DNA mutations related to adult mitochondrial disease. Ann. Neurol. 77, 753–759 (2015).

14. Gupta, R. et al. Nuclear genetic control of mtDNA copy number and heteroplasmy in humans. Nature 620, 839–848 (2023).

15. Suomalainen, A. & Battersby, B. J. Mitochondrial diseases: the contribution of organelle stress responses to pathology. Nat. Rev. Mol. Cell Biol. 19, 77–92 (2018).

16. Kang, Y. et al. Ancestral allele of DNA polymerase gamma modifies antiviral tolerance. Nature 628, 844–853 (2024).

17. Warren, E. B. et al. Inflammatory and interferon gene expression signatures in patients with mitochondrial disease. J. Transl. Med. 21, 331 (2023).

18. Primiano, G. et al. Inflammatory profile in mitochondrial diseases: A cohort study. Eur. J. Neurol. 30, 3409–3410 (2023).

19. Walker, M. A. et al. Predisposition to infection and SIRS in mitochondrial disorders: 8 years’ experience in an academic center. J. Allergy Clin. Immunol. Pract. 2, 465–468, 468.e1 (2014).

20. Eom, S. et al. Cause of Death in Children With Mitochondrial Diseases. Pediatr. Neurol. 66, 82– 88 (2017).

21. Edmonds, J. L. et al. The otolaryngological manifestations of mitochondrial disease and the risk of neurodegeneration with infection. Arch. Otolaryngol. Head Neck Surg. 128, 355–362 (2002).

22. Murray, P. J. et al. Macrophage Activation and Polarization: Nomenclature and Experimental Guidelines. Immunity 41, 14–20 (2014).

23. Ryan, D. G., Peace, C. G. & Hooftman, A. Basic Mechanisms of Immunometabolites in Shaping the Immune Response. J. Innate Immun. 15, 925–943 (2023).

24. Seim, G. L. et al. Two-stage metabolic remodelling in macrophages in response to lipopolysaccharide and interferon-γ stimulation. Nat. Metab. 1, 731–742 (2019).

25. Mills, E. L., Kelly, B. & O’Neill, L. A. J. Mitochondria are the powerhouses of immunity. Nat. Immunol. 18, 488–498 (2017).

26. Wang, Y., Li, N., Zhang, X. & Horng, T. Mitochondrial metabolism regulates macrophage biology. J. Biol. Chem. 297, 100904 (2021).

27. Cai, S. et al. Mitochondrial dysfunction in macrophages promotes inflammation and suppresses repair after myocardial infarction. J. Clin. Invest. 133, e159498 (2023).

28. Trifunovic, A. et al. Premature ageing in mice expressing defective mitochondrial DNA polymerase. Nature 429, 417–423 (2004).

29. Hooftman, A. et al. Macrophage fumarate hydratase restrains mtRNA-mediated interferon production. Nature 615, 490–498 (2023).

30. O’Carroll, S. M. et al. Itaconate drives mtRNA-mediated type I interferon production through inhibition of succinate dehydrogenase. Nat. Metab. (2024) doi:10.1038/s42255-024-01145-1.

31. Tang, J. et al. Disruption of glucose homeostasis by bacterial infection orchestrates host innate immunity through NAD+/NADH balance. Cell Rep. 43, (2024).

32. McNab, F., Mayer-Barber, K., Sher, A., Wack, A. & O’Garra, A. Type I interferons in infectious disease. Nat. Rev. Immunol. 15, 87–103 (2015).

33. Zecchini, V. et al. Fumarate induces vesicular release of mtDNA to drive innate immunity. Nature 615, 499–506 (2023).

34. Tábara, L.-C., Segawa, M. & Prudent, J. Molecular mechanisms of mitochondrial dynamics. Nat. Rev. Mol. Cell Biol. (2024) doi:10.1038/s41580-024-00785-1.

35. D, T., et al. SLP-2 is required for stress-induced mitochondrial hyperfusion. EMBO J. 28, (2009).

36. He, B. et al. Mitochondrial cristae architecture protects against mtDNA release and inflammation. Cell Rep. 41, 111774 (2022).

37. McShane, E. et al. A kinetic dichotomy between mitochondrial and nuclear gene expression processes. Mol. Cell 84, 1541–1555.e11 (2024).

38. Palmieri, E. M. et al. Nitric oxide orchestrates metabolic rewiring in M1 macrophages by targeting aconitase 2 and pyruvate dehydrogenase. Nat. Commun. 11, 698 (2020).

39. Jha, A. K. et al. Network Integration of Parallel Metabolic and Transcriptional Data Reveals Metabolic Modules that Regulate Macrophage Polarization. Immunity 42, 419–430 (2015).

40. Lampropoulou, V. et al. Itaconate Links Inhibition of Succinate Dehydrogenase with Macrophage Metabolic Remodeling and Regulation of Inflammation. Cell Metab. 24, 158–166 (2016).

41. Reynolds, M. B. et al. Type I interferon governs immunometabolic checkpoints that coordinate inflammation during Staphylococcal infection. Cell Rep. 43, 114607 (2024).

42. Birsoy, K. et al. An Essential Role of the Mitochondrial Electron Transport Chain in Cell Proliferation Is to Enable Aspartate Synthesis. Cell 162, 540–551 (2015).

43. Sullivan, L. B. et al. Supporting Aspartate Biosynthesis Is an Essential Function of Respiration in Proliferating Cells. Cell 162, 552–563 (2015).

44. Liu, P.-S. et al. α-ketoglutarate orchestrates macrophage activation through metabolic and epigenetic reprogramming. Nat. Immunol. 18, 985–994 (2017).

45. Martínez-Reyes, I. & Chandel, N. S. Mitochondrial TCA cycle metabolites control physiology and disease. Nat. Commun. 11, 102 (2020).

46. Mullen, A. R. et al. Reductive carboxylation supports growth in tumour cells with defective mitochondria. Nature 481, 385–388 (2012).

47. Metallo, C. M. et al. Reductive glutamine metabolism by IDH1 mediates lipogenesis under hypoxia. Nature 481, 380–384 (2011).

48. Burr, S. P. et al. Mitochondrial Protein Lipoylation and the 2-Oxoglutarate Dehydrogenase Complex Controls HIF1α Stability in Aerobic Conditions. Cell Metab. 24, 740–752 (2016).

49. Heinz, A. et al. Itaconate controls its own synthesis via feedback-inhibition of reverse TCA cycle activity at IDH2. Biochim. Biophys. Acta Mol. Basis Dis. 1868, 166530 (2022).

50. Jacobs, A. T. & Ignarro, L. J. Lipopolysaccharide-induced expression of interferon-beta mediates the timing of inducible nitric-oxide synthase induction in RAW 264.7 macrophages. J. Biol. Chem. 276, 47950–47957 (2001).

51. Palmieri, E. M. et al. Pyruvate dehydrogenase operates as an intramolecular nitroxyl generator during macrophage metabolic reprogramming. Nat. Commun. 14, 5114 (2023).

52. Van den Bossche, J., et al. Mitochondrial Dysfunction Prevents Repolarization of Inflammatory Macrophages. Cell Rep. 17, 684–696 (2016).

53. Tannahill, G. M. et al. Succinate is an inflammatory signal that induces IL-1β through HIF-1α. Nature 496, 238–242 (2013).

54. Ryan, D. G. et al. Disruption of the TCA cycle reveals an ATF4-dependent integration of redox and amino acid metabolism. eLife https://elifesciences.org/articles/72593 (2021) doi:10.7554/eLife.72593.

55. Shi, W., Cassmann, T. J., Bhagwate, A. V., Hitosugi, T. & Ip, W. K. E. Lactic acid induces transcriptional repression of macrophage inflammatory response via histone acetylation. Cell Rep. 43, (2024).

56. Timblin, G. A. et al. Mitohormesis reprogrammes macrophage metabolism to enforce tolerance. Nat. Metab. 3, 618–635 (2021).

57. Kruk, S. K. et al. Vulnerability of pediatric patients with mitochondrial disease to vaccine-preventable diseases. J. Allergy Clin. Immunol. Pract. 7, 2415–2418.e3 (2019).

58. Wculek, S. K. et al. Oxidative phosphorylation selectively orchestrates tissue macrophage homeostasis. Immunity 56, 516–530.e9 (2023).

59. Zhang, J., et al. Antigen receptor stimulation induces purifying selection against pathogenic mitochondrial tRNA mutations. JCI Insight 8, e167656 (2023).

60. Karaghiosoff, M. et al. Central role for type I interferons and Tyk2 in lipopolysaccharide-induced endotoxin shock. Nat. Immunol. 4, 471–477 (2003).

61. Mahieu, T. et al. The wild-derived inbred mouse strain SPRET/Ei is resistant to LPS and defective in IFN-beta production. Proc. Natl. Acad. Sci. U. S. A. 103, 2292–2297 (2006).

62. Crow, Y. J. & Stetson, D. B. The type I interferonopathies: 10 years on. Nat. Rev. Immunol. 22, 471–483 (2022).

63. Keshavan, N., Mhaldien, L., Gilmour, K. & Rahman, S. Interferon Stimulated Gene Expression Is a Biomarker for Primary Mitochondrial Disease. Ann. Neurol. 96, 1185–1200 (2024).

64. McGlasson, S., Jury, A., Jackson, A. & Hunt, D. Type I interferon dysregulation and neurological disease. Nat. Rev. Neurol. 11, 515–523 (2015).

65. Zhang, W. et al. Lactate Is a Natural Suppressor of RLR Signaling by Targeting MAVS. Cell 178, 176–189.e15 (2019).

66. Li, H. et al. AARS1 and AARS2 sense l-lactate to regulate cGAS as global lysine lactyltransferases. Nature 634, 1229–1237 (2024).

67. Malik, A. N., Czajka, A. & Cunningham, P. Accurate quantification of mouse mitochondrial DNA without co-amplification of nuclear mitochondrial insertion sequences. Mitochondrion 29, 59–64 (2016).

68. Zhong, Z. et al. New mitochondrial DNA synthesis enables NLRP3 inflammasome activation. Nature 560, 198–203 (2018).

69. Pearce, S. F. et al. Maturation of selected human mitochondrial tRNAs requires deadenylation. eLife 6, e27596 (2017).

70. Lebeau, J. et al. The PERK Arm of the Unfolded Protein Response Regulates Mitochondrial Morphology during Acute Endoplasmic Reticulum Stress. Cell Rep. 22, 2827–2836 (2018).

71. Dagda, R. K. et al. Loss of PINK1 Function Promotes Mitophagy through Effects on Oxidative Stress and Mitochondrial Fission *. J. Biol. Chem. 284, 13843–13855 (2009).

72. Shah, A. D., Goode, R. J. A., Huang, C., Powell, D. R. & Schittenhelm, R. B. LFQ-Analyst: An Easy-To-Use Interactive Web Platform To Analyze and Visualize Label-Free Proteomics Data Preprocessed with MaxQuant. J. Proteome Res. 19, 204–211 (2020).

73. Cader, M. Z. et al. FAMIN Is a Multifunctional Purine Enzyme Enabling the Purine Nucleotide Cycle. Cell 180, 278–295.e23 (2020).

74. Misheva, M. et al. Oxylipin metabolism is controlled by mitochondrial β-oxidation during bacterial inflammation. Nat. Commun. 13, 139 (2022).

75. Cader, M. Z. et al. C13orf31 (FAMIN) is a central regulator of immunometabolic function. Nat. Immunol. 17, 1046–1056 (2016).

76. Ge, S. X., Jung, D. & Yao, R. ShinyGO: a graphical gene-set enrichment tool for animals and plants. Bioinformatics 36, 2628 (2019).

77. Subramanian, A. et al. Gene set enrichment analysis: a knowledge-based approach for interpreting genome-wide expression profiles. Proc. Natl. Acad. Sci. U. S. A. 102, 15545–15550 (2005).

